# Sex–Specific Remodeling Phenotypes of the Tricuspid Valve Leaflets in an Ovine Model of Functional Tricuspid Regurgitation

**DOI:** 10.64898/2026.08.05.743037

**Authors:** Colton J. Kostelnik, Magda L. Piekarska, Shreya Sreedhar, Chien-Yu Lin, Aarya Shah, Boguslaw Gaweda, Austin J. Goodyke, Yujun Xu, Kartik Balachandran, Layla Parast, Matthew R. Bersi, Tomasz A. Timek, Manuel K. Rausch

## Abstract

**Background:** Moderate to severe tricuspid regurgitation (TR) affects approximately 1.6 million Americans, yet more than 90% of patients with significant TR remain untreated. Women exhibit higher TR prevalence and more rapid disease progression than men, but the valve-intrinsic mechanisms underlying these sex disparities remain unclear. We hypothesized that sex and circulating testosterone influence tricuspid leaflet remodeling during right-sided pressure overload.

**Methods:** Female, castrated male (C-Male), and non-castrated male (NC-Male) adult Dorset sheep (n = 45) underwent pulmonary artery banding (PAB) and were followed for 13 ± 1.5 weeks. Tricuspid leaflets were evaluated using morphometry, 3D profilometry, biaxial mechanical testing, histology, and bulk RNA sequencing. Sex-stratified differential gene expression was performed, and pathway enrichment of key biological processes were compared between sexes.

**Results:** PAB produced a uniform hemodynamic stimulus and equivalent moderate-to-severe TR across sex groups. Despite similar TR burden, leaflet remodeling diverged substantially by sex and castration status. C-Males developed the broadest remodeling phenotype, characterized by diffuse multi-leaflet growth, thickening, increased nuclei count, and low-strain stiffening. Females demonstrated more restricted leaflet and region-specific structural and cellular changes, along with circumferential low-strain stiffening. NC-Males exhibited preferential septal remodeling characterized by growth, thickening, increased nuclei count, and radial high-strain stiffening. Transcriptomic analysis revealed that females upregulated a focused matricellular remodeling program enriched for extracellular space organization (67 DEGs; FDR=0.025), whereas C-Males activated coordinated extracellular matrix and apoptosis-regulatory programs (388 DEGs; FDR=0.009). In contrast, NC-Males exhibited broad transcriptional response (406 DEGs) without significant pathway enrichment.

**Conclusions:** Tricuspid leaflet maladaptation during pressure overload is sex-dependent and testosterone-sensitive, involving distinct structural, mechanical, and transcriptional remodeling programs. These findings identify sex and testosterone status as previously under-recognized modulators of tricuspid valve remodeling and may help explain clinical sex disparities in TR progression.

**NOVELTY AND SIGNIFICANCE:** *What is known?:*

- Pulmonary hypertension and right ventricular pressure overload are linked to tricuspid leaflet remodeling through leaflet thickening, enlargement, and altered mechanical properties.
- Sex and sex-steroid hormones regulate fibrosis and extracellular matrix remodeling in cardiovascular tissues, but their role in tricuspid leaflet remodeling remains poorly understood.

*What new information does this article contribute?:*

- Sex and circulating testosterone status influence the magnitude, spatial distribution, biomechanical behavior, and transcriptional organization of tricuspid leaflet remodeling during pressure overload.
- Females, castrated males, and non-castrated males develop distinct remodeling programs characterized by focused matricellular remodeling, coordinated extracellular matrix/apoptosis signaling, and diffuse transcriptional activation, respectively.
- These findings identify sex and hormonal status as biological regulators of tricuspid valve maladaptation during functional tricuspid regurgitation.

*Summary:* Sex differences in tricuspid regurgitation progression are recognized clinically, yet the mechanobiological basis underlying these disparities remains poorly understood. Using a controlled ovine model of pressure overload–induced secondary tricuspid regurgitation, we demonstrated that tricuspid leaflet maladaptation is a sex-specific and testosterone-sensitive process spanning structural, mechanical, and transcriptional scales. Under comparable hemodynamic overload, all animals developed significant tricuspid regurgitation, but leaflet remodeling patterns diverged substantially across sexes. Castrated male sheep exhibited the broadest maladaptive phenotype, characterized by diffuse multi-leaflet growth and thickening, increased low-stretch stiffness, and coordinated extracellular matrix and apoptosis-regulatory transcriptional programs. Female sheep developed more spatially restricted remodeling accompanied by a focused matricellular and extracellular matrix secretory response, whereas non-castrated male sheep demonstrated selective leaflet remodeling with broad, but less coordinated, transcriptional activation. Different remodeling patterns emerged in females and castrated males despite comparable testosterone levels, suggesting that testosterone depletion alone does not fully explain these tricuspid valve remodeling phenotypes. These findings establish sex and testosterone status as previously underrecognized biological regulators of tricuspid leaflet maladaptation and support the emerging view that valve leaflets are active, mechanobiologically responsive, participants in functional tricuspid regurgitation progression.

## 1. INTRODUCTION

Tricuspid valve regurgitation (TR) is increasingly recognized as a major public health concern. Approximately 1.6 million US residents are diagnosed with moderate-to-severe TR, yet more than 90% of patients remain untreated [1,2].. This treatment gap reflects both the clinical complexity of right-sided heart disease and our incomplete understanding of disease progression. Epidemiological studies have established sex as a major determinant of TR prevalence and clinical presentation. Female patients often exhibit higher prevalence in TR prevalence, delayed presentation, and worse outcomes than male patients [2–5]. Despite these well-established sex differences, the biological mechanisms by which sex influences valve remodeling and TR progression remain poorly understood [6].

Functional or secondary TR has historically been viewed as a consequence of valve-extrinsic factors such as right ventricular (RV) dilation, annular enlargement, and papillary muscle displacement [2,7,8]. However, tricuspid valve leaflets are now recognized as active, mechanosensitive tissues that grow and remodel in response to altered mechanical loading [9–11]. Our prior ovine secondary TR studies have shown that tricuspid leaflets undergo maladaptive remodeling in response to either biventricular heart failure or isolated right ventricular (RV) pressure overload, characterized by leaflet enlargement, thickening, stiffening, and extracellular matrix (ECM) reorganization [12–15]. These remodeling responses are spatially heterogeneous and likely driven by differences in the magnitude and distribution of in vivo areal strains and intrinsic ECM properties [16–18]. Increased leaflet thickness and stiffness significantly reduce coaptation area, indicating that leaflet remodeling contributes to valve dysfunction rather than being a consequence of it [19]. Thus, the tricuspid valve itself is not an innocent bystander but a direct contributor to disease progression [20].However, the computational models informing these findings were parameterized using remodeling data from our previous male-only experimental studies, leaving two major questions unaddressed: whether leaflet remodeling itself differs by sex, and whether any such difference is driven by sex hormone signaling.

Sex steroid hormones are key regulators of cardiac tissue remodeling, acting on resident fibroblasts and directing ECM dynamics [21]. For example, in vitro studies have shown that physiological testosterone levels in male rodents suppressed TGF-β and angiotensin II-dependent profibrotic signaling, myofibroblast differentiation, collagen synthesis, and cell proliferation in cardiac fibroblasts [22,23]. In clinical studies, testosterone deficiency in men has been associated with advanced coronary disease, a worse functional status, and higher rates of major adverse cardiovascular events and mortality [24,25]. Similarly, in vivo studies have shown that estradiol attenuates RV fibrosis, collagen accumulation, and proapoptotic signaling in response to pressure overload [26,27]. However, estrogen’s role in cardiac remodeling is notably dimorphic, promoting favorable myocardial adaptation while producing inconsistent, model-dependent effects on the vasculature [28]. These findings support the concept that sex hormone-dependent pathways operate in cardiac cells and tissues and play a significant role in cardiac remodeling [29]. However, it remains unknown whether these mechanisms extend to tricuspid valve leaflets and contribute to maladaptation in secondary TR.

To determine the roles of biological sex and circulating testosterone in tricuspid leaflet remodeling, we leveraged a well-established ovine pulmonary artery banding (PAB) model to induce pulmonary hypertension, RV remodeling, secondary TR, and the associated tricuspid leaflet remodeling in Female, castrated male (C-Male), and non-castrated male (NC-Male) sheep [12,30]. We then assessed leaflet structure, mechanics, and transcriptomics to fully characterize remodeling phenotypes. Based on the profibrotic consequences of testosterone deficiency in males, we hypothesized that C-Males would exhibit the most pronounced and diffuse maladaptive remodeling response, while Females would display more targeted adaptations and NC-Males would develop an intermediate remodeling phenotype.

## 2. MATERIALS & METHODS

### 2.1 Animal Studies

Healthy adult Dorset sheep were assigned to control (CTL) or pulmonary artery banding (PAB) treatment groups (**Table 1**). All procedures were approved by the Michigan State University Institutional Animal Care and Use Committee (protocols PROTO202200120, PROTO202500129). PAB animals were followed for 12-15 weeks after banding to monitor RV pressure overload and remodeling, while CTL animals did not undergo banding or sham surgery. At the terminal timepoint, we collected animal blood plasma to quantify concentrations of cytokine and sex steroid hormones via multiplex immunoassay. We excised the tricuspid valves from all animals for structural, mechanical, and transcriptomic characterization.

**Table 1.** Terminal hemodynamics and echocardiographic variables in control (CTL) and pulmonary artery banding (PAB) sheep stratified by sex and group.

|  | Female |  | Castrated Male<br>(C-Male) |  | Non-castrated Male<br>(NC-Male) |  |
| --- | --- | --- | --- | --- | --- | --- |
|  | CTL<br>(n = 9) | PAB<br>(n = 10) | CTL<br>(n = 7) | PAB<br>(n = 9) | CTL<br>(n = 5) | PAB<br>(n = 5) |
| *Hemodynamic Data* |  |  |  |  |  |  |
| sPAP, mmHg | 16.0 ± 2.6 | 27.1 ± 4.7 | 14.6 ± 5.3 | 31.0 (17.75)<sup>*</sup> | 14.8 ± 3.7 | 36.6 ± 7.5 |
| mPAP, mmHg | 10.0 ± 4.2<sup>#,§</sup> | 17.1 ± 3.6 | 8.3 ± 4.1 | 20.9 (16.4)<sup>*,¥</sup> | 9.4 ± 2.7 | 19.8 ± 2.3 |
| dPAP, mmHg | 8.0 (7.5)<sup>#,§</sup> | 9.0 ± 4.3<sup>*</sup> | 5.0 ± 3.7 | 11.5 (18.3)<sup>*,¥</sup> | 6.6 ± 2.1 | 6.2 ± 0.8 |
| sAP, mmHg | 99.6 ± 7.6 | 88.2 ± 12.8<sup>§</sup> | 85.9 ± 6.3 | 77.9 ± 25.6<sup>*,¥</sup> | 82.8 ± 10.0 | 98.2 ± 19.7<sup>*</sup> |
| mAP, mmHg | 86.1 ± 11.1 | 76.5 ± 12.5<sup>§</sup> | 71.6 ± 9.9 | 66.4 ± 27.0<sup>¥</sup> | 70.6 ± 10.2 | 86.6 ± 16.8<sup>*</sup> |
| dAP, mmHg | 74.7 ± 12.1 | 67.1 ± 12.5<sup>§</sup> | 61.0 ± 12.45 | 57.8 ± 30.0<sup>¥</sup> | 62.0 ± 12.3 | 77.8 ± 16.6<sup>*</sup> |
| *Echocardiographic Data* |  |  |  |  |  |  |
| TR (0-4) | 1.0 (0.0) | 4.0 (1.8)<sup>*</sup> | 1.0 (1.0) | 3.5 (1)<sup>*</sup> | 1.4 ± 0.4 | 2.6 ± 1.3<sup>*</sup> |
| RVFAC (%) | 61.8 ± 8.7 | 49.8 ± 9.5 | 64.6 ± 9.4 | 38.5 ± 3.7<sup>*</sup> | 62.8 ± 13.4 | 45.0 ± 19.8<sup>*</sup> |
| TVAD<sub>idx</sub> (cm/kg) | 0.04 ± 0.01 | 0.05 ± 0.01<sup>*</sup> | 0.04 ± 0.01 | 0.04 (0.01)<sup>*</sup> | 0.04 ± 0.01 | 0.05 ± 0.01<sup>*</sup> |
| RAA<sub>idx</sub> (cm<sup>2</sup>/kg) | 0.12 ± 0.04 | 0.20 ± 0.05<sup>*</sup> | 0.14 ± 0.03 | 0.16 (0.04) | 0.12 ± 0.01 | 0.18 ± 0.06 |
| RVDd1<sub>idx</sub> (cm/kg) | 0.04 ± 0.01 | 0.05 ± 0.01<sup>*</sup> | 0.05 ± 0.01 | 0.05 ± 0.01 | 0.04 ± 0.01 | 0.05 ± 0.01 |
| RVDd2<sub>idx</sub> (cm/kg) | 0.03 ± 0.01 | 0.04 ± 0.01<sup>*</sup> | 0.03 (0.01) | 0.03 ± 0.01 | 0.02 ± 0.01 | 0.03 ± 0.01 |
| RVDd3<sub>idx</sub> (cm/kg) | 0.07 ± 0.01 | 0.08 ± 0.01 | 0.07 ± 0.01 | 0.07 ± 0.01 | 0.06 ± 0.01 | 0.08 ± 0.01 |
| *Plasma Hormone Data* |  |  |  |  |  |  |
| Testosterone (ng/mL) | 0.03 ± 0.02<sup>§</sup> | 0.02 (0.03)<sup>§</sup> | 0.03 ± 0.02<sup>¥</sup> | 0.02 (0.01)<sup>¥</sup> | 2.1 ± 2.19 | 1.3 ± 1.15 |
| Progesterone (ng/mL) | 0.3 ± 0.23 | 0.2 (0.91) | 0.1 ± 0.05 | 0.10 ± 0.09 | 0.12 ± 0.02 | 0.04 (0.10) |
| Estradiol (ng/mL) | 0.08 ± 0.03 | 0.03 (0.05) | 0.04 (0.03) | 0.03 ± 0.02 | 0.04 ± 0.02 | 0.05 ± 0.04 |
| Cortisol (ng/mL) | 40.3 ± 12.33 | 44.3 ± 12.95 | 37.3 ± 12.7 | 41.7 ± 18.5 | 33.3 ± 6.43 | 41.4 ± 13.21 |

Values are mean ± standard deviation or median (interquartile range) as determined by the Shapiro-Wilk normality test. Statistical differences were determined from pairwise comparisons of linear mixed effect model.
CTL = control, PAB = pulmonary artery banding, PAP = pulmonary artery pressure, AP = arterial pressure, s = systolic, d = diastolic, m = mean, TR = tricuspid regurgitation, FAC = fractional area change, TVAD = tricuspid valve annular diameter, idx = indexed by weight, RAA = right atrium area, RVD = right ventricle diameter.
\* p < 0.05 (CTL vs. PAB), # p < 0.05 (Female vs. C-Male), ¥ p < 0.05 (C-Male vs. NC-Male), § p < 0.05 (Female vs. NC-Male)

### 2.2 Tricuspid Leaflet Characterization

Tricuspid valve leaflets were assessed for morphology, thickness, biaxial mechanical behavior, histological cell density, and transcriptomics to characterize treatment- and sex-dependent remodeling to RV pressure overload-induced TR. Detailed methods for all experiments are provided in the Supplemental Materials

### 2.3 Statistical Analysis

Forty-five sheep were used in this study and were age-matched across sexes within each group. We analyzed morphological, thickness, biaxial, and histological data using linear mixed-effects models in R (lme4 package), with group, sex, and leaflet as fixed effects and subject as a random effect. We used Type III ANOVA with Satterthwaite’s approximation to assess significance of main effects and interactions. We conducted post-hoc pairwise comparisons using estimated marginal means (emmeans package) with false discovery rate (FDR) correction where applicable. We defined statistical significance as p < 0.05 and report data as mean ± standard deviation.

## 3. RESULTS

### 3.1 PAB induces sex–dependent RV remodeling in a pressure–overload TR model

PAB acutely elevated systolic and mean pulmonary artery pressures and worsened TR grade in all sex groups, confirming an isolated right-heart pressure overload stimulus (**Table S2**). At 13 ± 1.5 weeks, all PAB animals developed moderate-to-severe TR with significant annular dilation (**Table 1**). However, different sex groups exhibited distinct RV changes. Female sheep exhibited geometric RV remodeling with enlarged right atrial area (70% increase, p = 0.02) and increased RV diameters (15% increase in basal diameter [d1], p = 0.04; 29% increase in mid-cavity diameter [d2], p = 0.003). In contrast, male sheep preserved chamber dimensions but developed impaired RV systolic function, with fractional area change decreasing by 26% in PAB C-Males (p < 0.001) and by 18% in PAB NC-Males (p = 0.04) (**Table 1**). At the terminal timepoint, only C-Males sustained mean and systolic pulmonary artery pressures significantly higher than their CTL counterparts (**Table 1**).

Castration effectively reduced circulating testosterone concentrations in C-Males to Female levels, while testosterone concentrations in NC-Males maintained significantly higher concentrations, approximately 40-100 times greater than in Females and C-Males (**Table 1**). Progesterone, estradiol, and cortisol showed no statistically significant differences across groups or sexes. PAB did not trigger a systemic pro-inflammatory cytokine response. Terminal plasma cytokine concentrations of IL-1β, IL-6, IL-8, IL-10, and TNF-α showed modest decreases, or no change, across sexes in the PAB group. Significant differences included lower IL–6, IL–8, and TNF-α in PAB NC–Males relative to CTL and higher IL–8 in PAB Females relative to PAB NC-Males (**Figure S1**).

Together, these data establish a pressure-driven TR model with minimal systemic inflammation and sex-dependent RV remodeling in the setting of effective testosterone deprivation in C-Males.

### 3.2 PAB drives sex- and leaflet-dependent growth

Given known sex disparities in TR prevalence and progression, we hypothesized that PAB would enlarge tricuspid leaflets in a sex- and leaflet-dependent manner. At baseline (CTL), Female anterior leaflets were smaller than NC-Male anterior leaflets (p = 0.04; **Figure 1c**), and NC-Male posterior leaflets were smaller than C-Male posterior leaflets (p = 0.03; **Figure 1d**), while septal leaflet size did not differ were observed between sex groups (**Figure 1b**). PAB reshaped, but did not eliminate these differences. Among PAB animals, female anterior and posterior leaflets remained smaller than C-Male leaflets (p = 0.01 and p = 0.002, respectively; **Figure 1a,c,d**). To assess global remodeling, we summed leaflet areas within each valve and observed that total leaflet area increased with PAB in all sexes (22% increase; all p < 0.04; **Figure S2b**), indicating all animals exhibited net leaflet growth regardless of sex. Among PAB animals, total leaflet area in Females remained smaller than C-Males (p = 0.005; **Figure S2b**).

**Figure 1.**
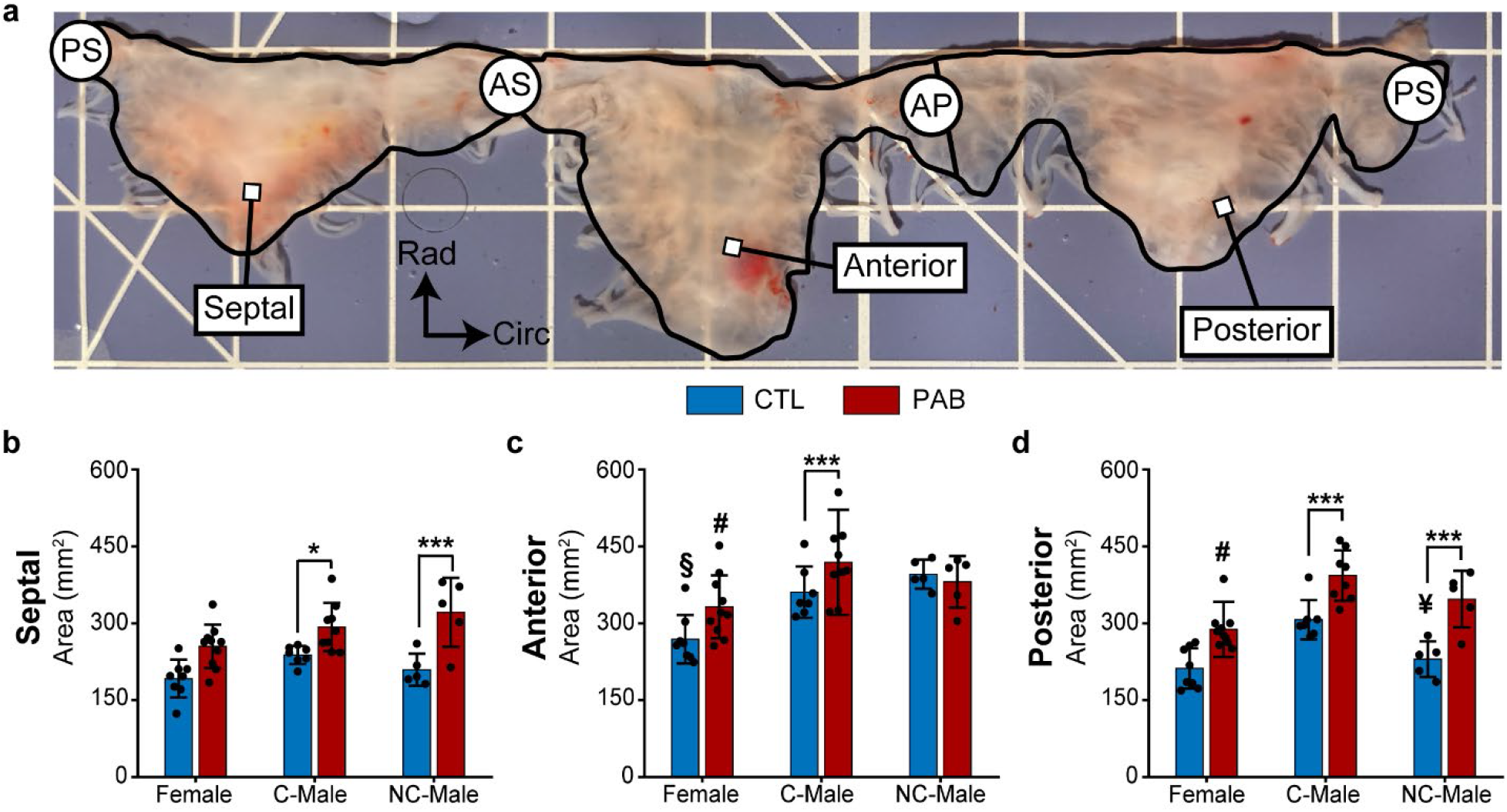
Pulmonary artery banding (PAB) enlarges tricuspid valve leaflet area with distinct patterns across leaflets and sex. (a) Representative tricuspid valve imaged on a calibrated grid (1-cm scale). Septal, anterior, and posterior leaflet boundaries are traced (black outlines), and commissures are labeled as posterior–septal (PS), antero–septal (AS), and antero–posterior (AP). Leaflet areas were measured between PS and AS for the septal leaflet, AS and AP for the anterior leaflet, and AP and PS for the posterior leaflet. (b–d) Leaflet area for the septal, anterior, and posterior leaflets, respectively, in control (CTL, blue) and PAB (red) sheep stratified by sex (Female, castrated male (C–Male), and non-castrated male (NC–Male)). Data are shown for individual animals with group means ± standard deviation. Statistical differences between groups were assessed using linear mixed–effects models with post hoc pairwise comparisons of estimated marginal means. Significant contrasts between groups (CTL vs. PAB) are denoted by (*) for p < 0.05 and (***) for p < 0.001. Significant contrasts between sexes are denoted by (#) for p <0.05 (C-Male vs. Female), (§) for p < 0.05 (NC-Male vs. Female), and (¥) for p < 0.05 (C-Male vs. NC-Male).

PAB induced leaflet-specific growth patterns that differed by sex (group *x* sex *x* leaflet interaction, p = 0.006; **Figure 1**). C-Males exhibited the largest remodeling response, with all three leaflets growing in area (anterior: +16%, posterior: +27%, septal: +23%; all p < 0.02; **Figure 1b–d**). NC-Males showed selective growth in the posterior and septal leaflets (50-52% increase, both p < 0.001; **Figure 1b,d**), whereas their anterior leaflets remained unchanged. Females displayed larger mean leaflet areas (8-15% increases) that did not reach statistical significance. Together, these findings demonstrate that PAB induces leaflet growth across all sexes, but leaflet specific remodeling patterns differ significantly based on sex and testosterone status. Testosterone deprived C-Males exhibited the most extensive and uniformly distributed leaflet enlargement, while Females showed more modest remodeling responses.

### 3.3 PAB induces regionally heterogeneous thickening with sex- and leaflet-specific patterns

Prior studies in ovine secondary TR models have linked leaflet thickening to impaired valve competence [15,18,19]. We first compared mean whole-leaflet thickness between groups stratified by sex. PAB induced leaflet thickening, but sex and the leaflet identity shaped the remodeling pattern (sex: p < 0.01; leaflet: p < 1.0E-10; **Figure 2**). At baseline (CTL), NC-Male anterior and posterior leaflets were thicker than C-Male leaflets (p = 0.003 and p = 0.04, respectively; **Figure 2c,d**); however, following PAB, female and C-Male leaflets thickened to levels comparable to NC-Male leaflets eliminating baseline sex differences. C-Males showed the largest remodeling response, with all three leaflets thickening with PAB (anterior: +41%, posterior: +48%, septal: +43%, all p < 0.001; **Figure 2b–d**). Females exhibited PAB-induced thickening limited to the septal (+22%, p = 0.035; **Figure 2b**) and posterior leaflets (+39%, p = 0.013; **Figure 2d**), whereas anterior leaflets did not change. NC-Males showed the smallest remodeling response to PAB with only the septal leaflets thickening (+27%, p = 0.015; **Figure 2b**). Representative thickness maps for CTL and PAB Female, C-Male, and NC-Male valves are shown in **Figures S3–S5**, respectively.

**Figure 2.**
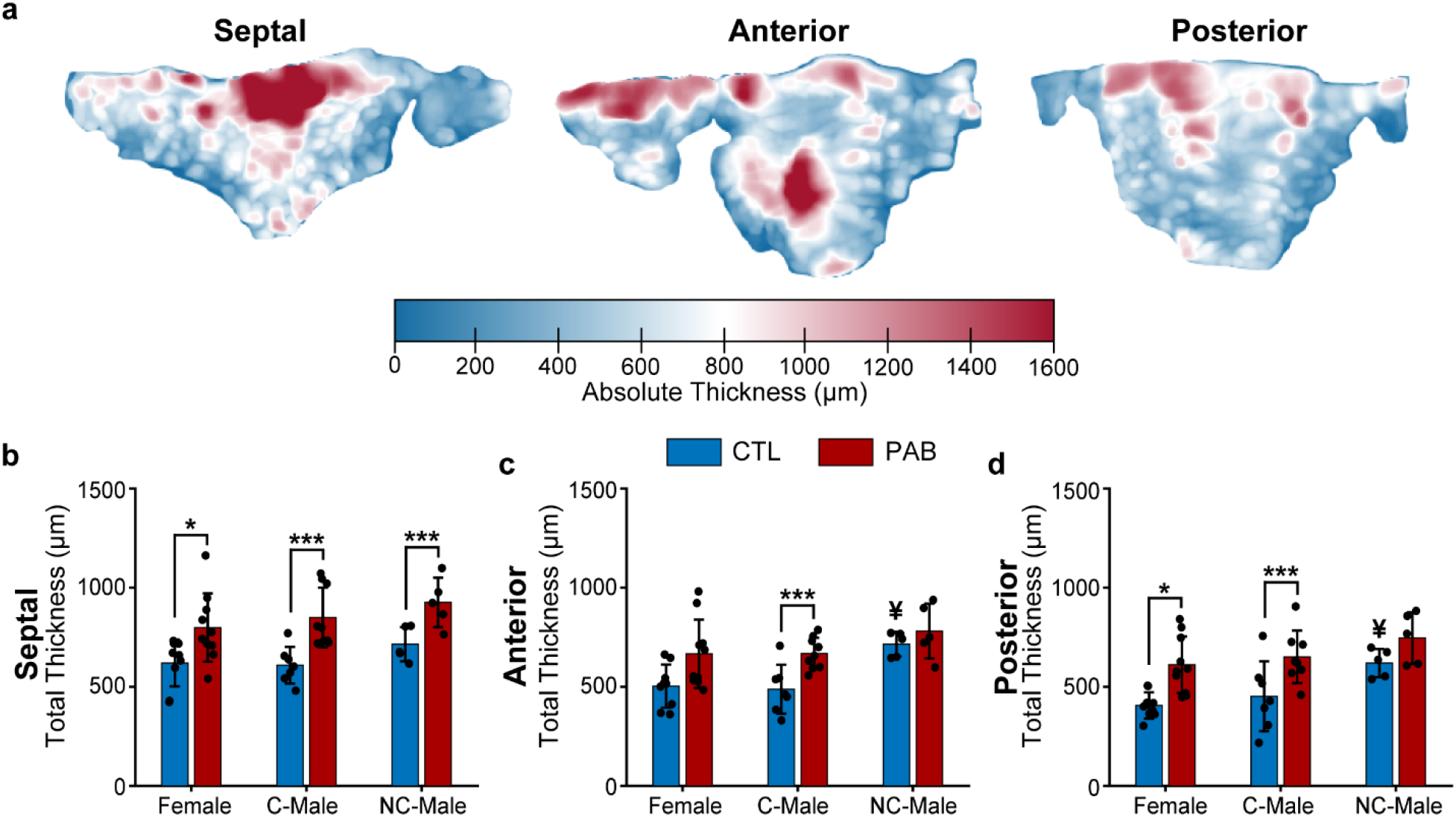
Pulmonary artery banding (PAB) induced leaflet thickening differs by leaflet and sex. (a) Representative profilometric thickness maps for a PAB septal, anterior, and posterior leaflet. Total leaflet thickness comparison between control (CTL, blue) and pulmonary artery banding (PAB, red) sheep stratified by sex (Female, castrated male (C–Male), and non-castrated male (NC–Male)). Total thickness of the (b) septal, (c) anterior, and (d) posterior leaflets was quantified from the profilometric thickness maps. Data are shown for individual animals with group means ± standard deviation. Statistical differences between groups were assessed using linear mixed–effects models with post hoc pairwise comparisons of estimated marginal means. Significant contrasts between treatment groups (CTL vs. PAB) are denoted by (*) for p < 0.05 and (***) for p < 0.001, while significant contrasts between sex groups are denoted by (¥) for p < 0.05 (C-Male vs. NC-Male).

We segmented leaflet thickness maps into a 3×3 regional grid (**Figure S6**) and found that PAB-induced thickening depended strongly on leaflet and location (group *x* radial position interaction, p < 0.0001; **Figure S7a-d**). C-Male leaflets exhibited the most extensive remodeling response, with significant thickening across nearly all leaflet regions. Female leaflets showed a more constrained pattern, with significant thickening concentrated in the near-annulus and free edge regions and belly thickening across the midline in all leaflet. NC-Males exhibited the most localized and leaflet-specific thickening response, with significant thickening largely confined to the near-annulus and free edge regions of the septal and posterior leaflets, and minimal thickening observed in the belly region or anterior leaflet. Together, these findings indicate that PAB drives regionally heterogeneous leaflet thickening across all sexes, while sex and testosterone status reshape the regional pattern of thickening rather than the overall magnitude.

### 3.4 PAB alters leaflet mechanical behavior in a sex- and leaflet-specific manner

Maladaptive tricuspid leaflet remodeling alters leaflet mechanical properties at low and high strain regimes [14,15,18]. We next asked whether these observed differences in sex-specific structural changes were accompanied by sex-specific alterations to leaflet mechanics. All leaflets exhibited the classic J-shaped tension–stretch response of collagenous soft tissues in both the circumferential (**Figure 3a–c**) and radial (**Figure 3g–i**) directions. PAB stiffened leaflets preferentially at low stretches (toe stiffness: group *x* direction interaction, p < 0.001, **Figure 3d,j**; transition stretch: group *x* direction interaction, p < 0.03, **Figure 3f,l**). Stiffness at high stretches (calf stiffness) depended on group *x* direction (p < 0.01; **Figure 3e,k**) but not on sex as a main effect.

**Figure 3.**
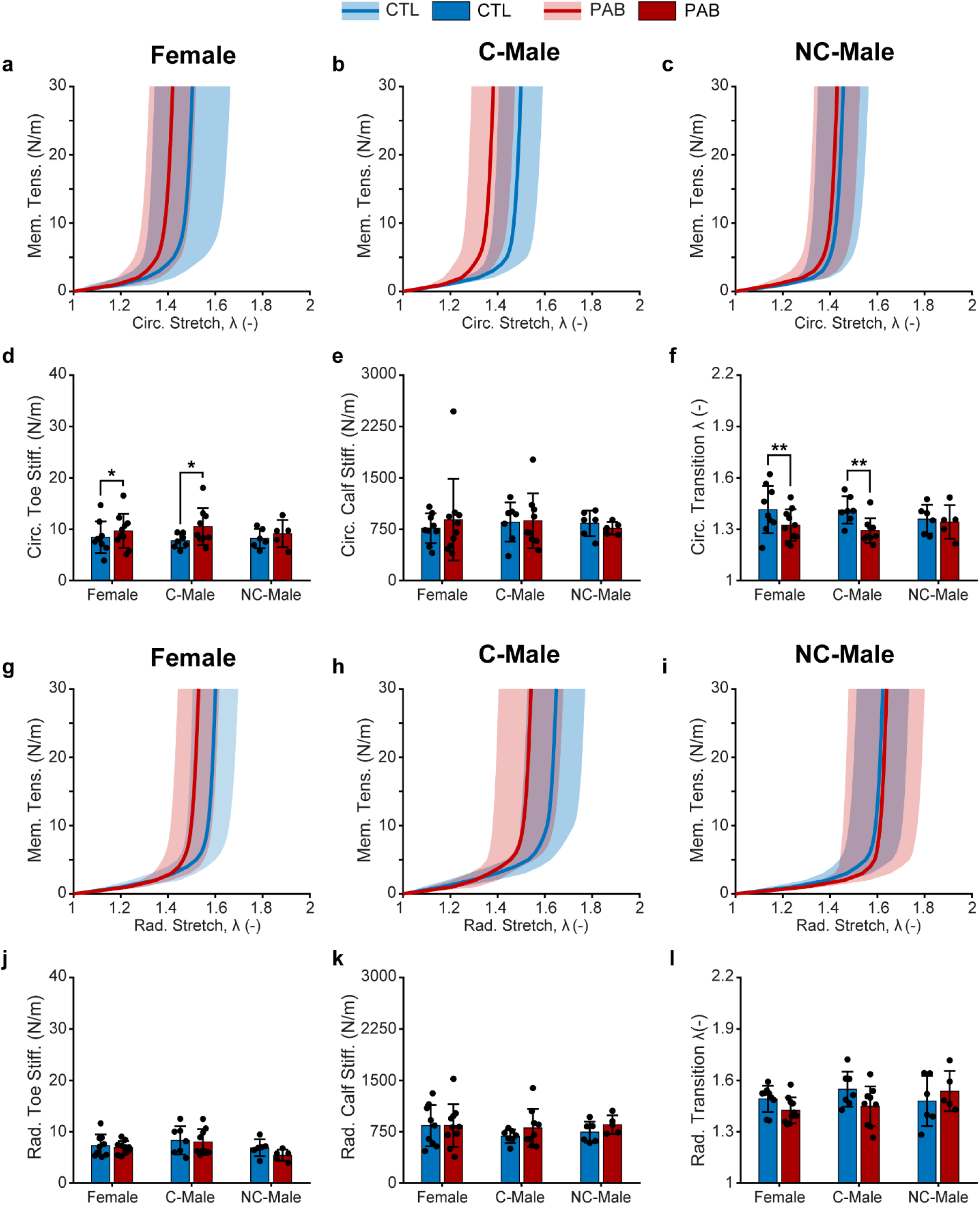
Pulmonary artery banding (PAB) alters the biaxial mechanical behavior of the anterior leaflet. Circumferential membrane tension (Mem. Tens.) and stretch curves for control (CTL, blue) and PAB (red) sheep are shown for (a) Female, (b) castrated male (C–Male), and (c) non–castrated male (NC–Male) sheep, with shaded regions denoting mean (solid line) and standard deviation (shaded). Corresponding circumferential mechanical metrics are presented as (d) toe stiffness, (e) calf stiffness, and (f) transition stretch (λ) for each sex group. Radial membrane tension–stretch curves for CTL and PAB sheep are shown for (g) Female, (h) C–Male, and (i) NC–Male sheep, with shaded regions denoting ± standard deviation, and the associated radial (j) toe stiffness, (k) calf stiffness, and (l) transition stretch (λ). Data are shown for individual animals with group means ± standard deviation. Statistical differences between groups were assessed using linear mixed–effects models with post hoc pairwise comparisons of estimated marginal means. Significant contrasts between groups (CTL vs. PAB) denoted by (*) for p < 0.05 and (**) for p < 0.01.

Consistent with prior work, the anterior leaflet showed the most remodeling [18]. Females and C-Males stiffened at low stretches, with circumferential toe stiffness increasing in both groups (females: +48%, C-Male: +45%; both p < 0.02; **Figure 3d**) and circumferential transition stretch decreasing (both - 10%; p < 0.01; **Figure 3f**), suggesting earlier collagen fiber engagement. Septal and posterior leaflets followed similar patterns (**Figures S9** and **S10**, respectively). Females and C-Males increased circumferential toe stiffness (both p < 0.01; **Figures S9d, S10d**), and female circumferential transition stretches decreased (all p < 0.01; **Figure 3f** and **Figures S9f, S10f**); the reduction in C-Male circumferential transition stretch was limited to the anterior leaflet. In contrast, NC-Males preserved low-stretch compliance, with circumferential toe stiffness and transition stretch remaining unchanged across all leaflets. Toe stiffness and transition stretch in the radial direction were unaffected by PAB across all leaflets (**Figure 3j,l** and **Figures S9j,l, S10j,l**).

PAB-induced high-stretch stiffening was only observed in NC-Male radial calf stiffness in the septal leaflet (+63%, p < 0.02; **Figure S9k**) and C-Male radial calf stiffness in the posterior leaflet (+60%, p < 0.02; **Figure S10k**). Female leaflets did not exhibit a significant change in high-stretch stiffness in any leaflet. Direct sex comparisons within the PAB group revealed additional posterior leaflet differences: NC-Males had lower circumferential toe stiffness than females and C-Males (p < 0.04; **Figure S10d**), and lower radial calf stiffness than C-Males (p < 0.04; **Figure S10k**). Collectively, these data show that female and C-Male leaflets undergo consistent circumferential stiffening at low stretch, whereas NC-Males preserve low-stretch compliance and exhibit more localized increases in high-stretch stiffness, particularly in the radial direction of the septal leaflet.

### 3.5 PAB induces sex–specific redistribution of leaflet nuclei density

Based on prior work linking cell nuclei density and matrix remodeling to leaflet maladaptation, we hypothesized that PAB would alter leaflet nuclei density in a sex- and leaflet-specific manner [15,18]. Mixed-effects analysis showed significant main effects of group and leaflet on total nuclei count (all p < 0.03; **Figure 4**), without significant interactions among these factors. PAB C-Males showed increased nuclei count in the anterior leaflet (+71%, p = 0.019 **Figure 4d**). PAB Females showed increased total nuclei in all leaflets but did not reach statistical significance (anterior: +60%; posterior: +67%; septal: +62%; p > 0.35; **Figure 4b**). NC-Males exhibited selective increases, with only septal leaflets containing more nuclei after PAB (+94%, p < 0.001; **Figure 4d**), which was significantly greater than in PAB females and C-Males (both p < 0.02; **Figure 4d**).

**Figure 4.**
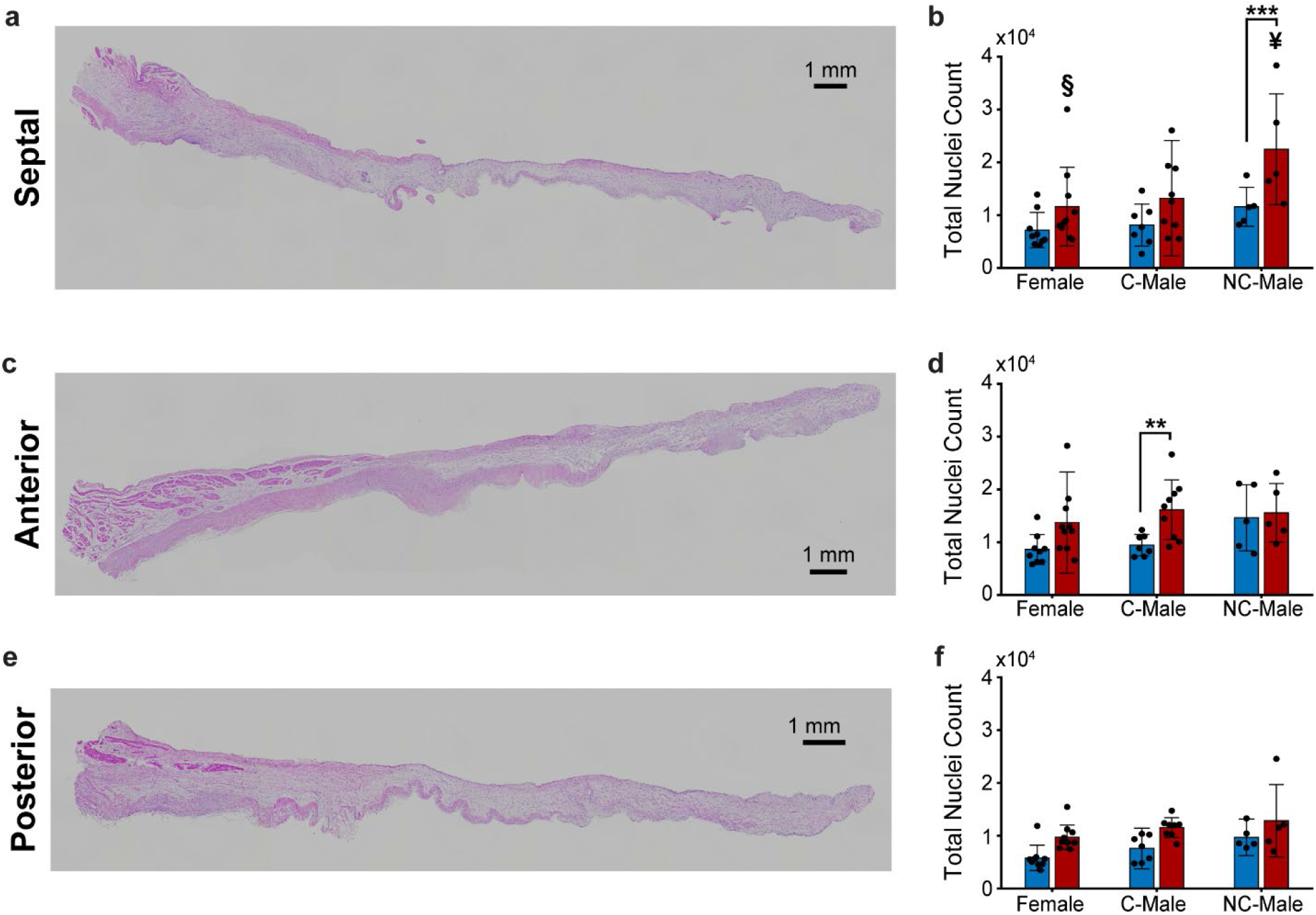
Total nuclei count increases in leaflets after pulmonary artery banding (PAB) in a sex-dependent manner. Representative H&E-stained cross-sectional histology images for (a) septal, (c) anterior, (e) posterior leaflets from a diseased (PAB) non-castrated male (NC-Male) sheep. Total cell nuclei comparison for (b) septal, (d) anterior, and (f) posterior leaflets from Female, castrated male (C-Male), and non-castrated male (NC-Male) sheep. Data are shown for individual animals with group means ± standard deviation. Statistical significance between groups (CTL vs. PAB) is denoted by (*) for p < 0.05, (**) for p < 0.01, and (***) for p < 0.001. Statistical significance between sexes is denoted by (§) for p < 0.05 (NC-Male vs. Female) and (¥) for p < 0.05 (C-Male vs. NC-Male).

Regional nuclei density varied by sex, leaflet, and radial position (all p < 0.01; **Figure 4** and **Figure S11**), with a significant sex *x* radial position interaction (p < 0.01). Significant regional increases in nuclei density were limited and largely confined to the ventricular surface at the free edge region. PAB Females showed a significant increase at the ventricular free edge of the anterior leaflet (**Figure S11c**), whereas PAB NC-Males showed a significant increase at the ventricular free edge of the septal and posterior leaflets (p < 0.05; **Figure S11b,d**). PAB C-Males showed no significant regional increases in nuclei density in any leaflet (**Figure S11c,d**). Together, these findings indicate that PAB modifies leaflet nuclei density in a sex- and leaflet-specific manner, with modest whole-leaflet increases and spatially restricted regional concentrated at the ventricular free edge region.

### 3.6 Sex and circulating testosterone shape ECM-focused transcriptional remodeling after PAB

We next asked whether PAB elicits sex-dependent transcriptional remodeling in the tricuspid valve. Using bulk RNA sequencing and a DESeq2 model that accounted for sex, group, and their interaction, we computed sex-stratified transcriptional changes. Principal component analysis revealed no clear separation of samples by leaflet or sex, and we therefore proceeded with a leaflet-pooled analytical approach all downstream analysis (**Figure S12a and S12b**). However, given the known differences in leaflet-specific remodeling, we performed sex- and leaflet-stratified differential expression analyses to directly test for sex- and leaflet-dependent transcriptional responses (**Figure S13**). At our predefined thresholds (adjusted p ≤ 0.05 and |log2(FC)| ≥ 0.5), all sexes exhibited some degree of differential gene expression, but the number of DEGs varied substantially by sex. Female sheep had the smallest DEG set (67 upregulated, 65 downregulated), while both male groups demonstrated larger transcriptional responses (C-Males: 388 upregulated, 354 downregulated; NC-Males: 406 upregulated, 219 downregulated; **Figure 5**). To assess the degree of transcriptional overlap, we compared upregulated and downregulated DEGs across sex groups (**Figure S14a and S14b**). The majority of DEGs were sex specific, with minimal overlap between groups in response to PAB. Notably, a small core of 11 genes were upregulated across all sex groups (e.g., LOXL2, CXCL14, TNC, METRNL, GLDN, and DAAM2), and 6 genes were downregulated across all sex groups (e.g., ADCYAP1R1, ERBB3, GAS1, IRF2BPL, SMTNL2, TGM2).

**Figure 5.**
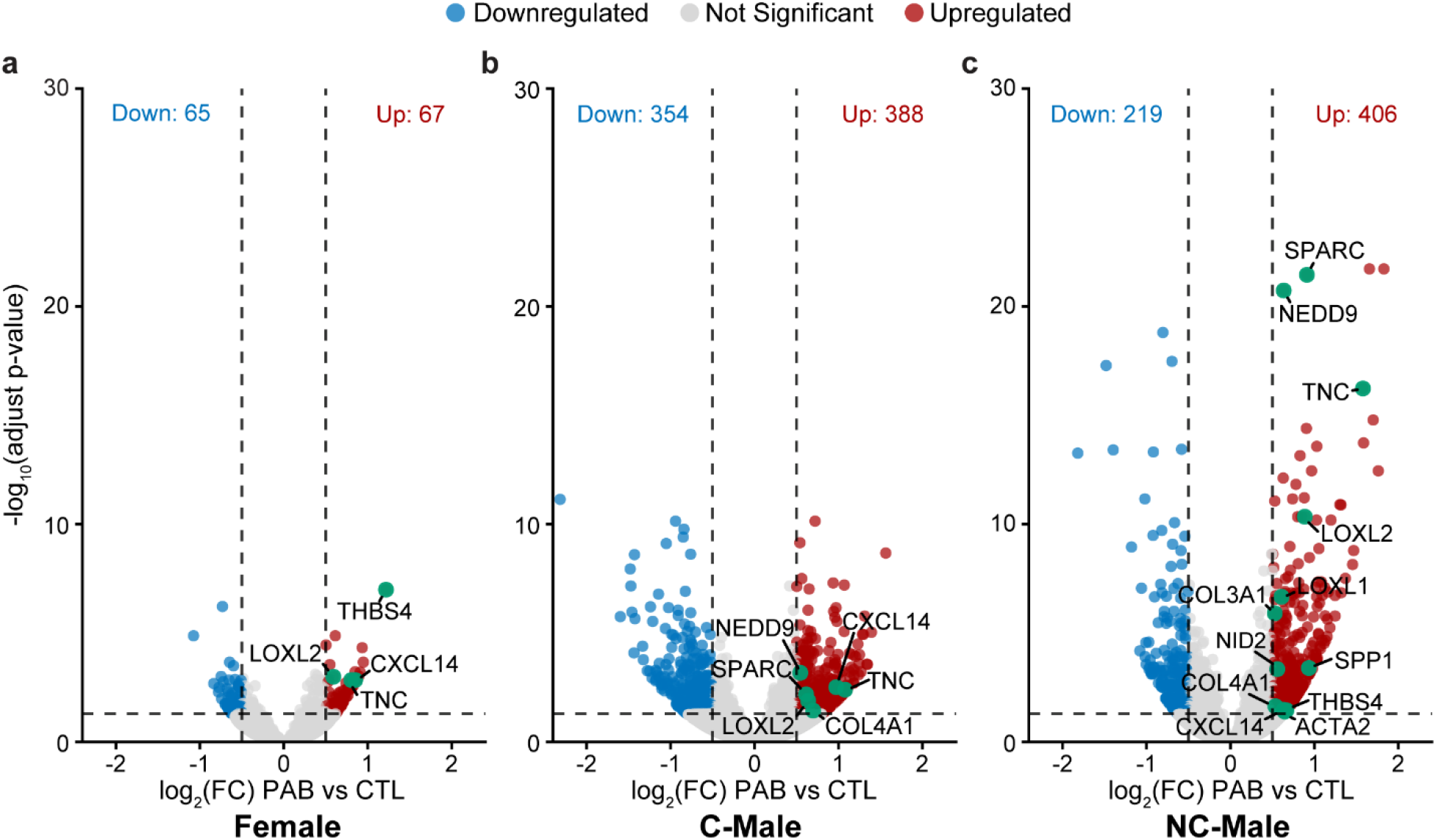
Sex–stratified differential gene expression in tricuspid valve leaflets after pulmonary artery banding (PAB). Volcano plots depict differential gene expression (PAB vs. control [CTL]) from bulk RNA-seq of tricuspid valve leaflets from (a) Female, (b) castrated male (C-Male), and (c) non-castrated male (NC-Male) sheep. Each point represents a gene, plotted by log2-fold change (FC) (PAB vs. CTL) on the x-axis and -log10(adjusted p-value) on the y-axis. Significantly upregulated and downregulated genes (adjusted p ≤ 0.05 and |log2(FC)| ≥ 0.5) are shown in red and blue, respectively; non-significant genes are shown in grey. Dashed lines indicate significance thresholds. The total number of differentially expressed genes is reported for each group. Selected extracellular matrix and remodeling-associated genes in each group are annotated.

Despite this shared activation of remodeling-associated transcription, the composition and organization of the response differed by sex and testosterone status. Female leaflets upregulated a compact ECM-secretory program, including matricellular and chemokine-associated genes such as THBS4, CXCL14, LOXL2, and TNC, consistent with ECM modification rather than broad matrix turnover. C-Males leaflets exhibited a different expression profile characterized by induction of both ECM and signaling-associated genes, including CXCL14, NEDD9, COL4A1, and ADAMTS family proteases (ADAMTS4, ADAMTS9, ADAMTSL2). In contrast, NC-Males leaflets showed upregulation of ECM structural, matricellular, and activation-associated genes, including SPARC, LOXL1/2, SPP1, ACTA2, and multiple collagen isoforms (COL1A2, COL3A1, COL4A1).

To further characterize these sex- and testosterone-associated remodeling programs, we examined a curated panel of immune, ECM/matricellular, matrix-modifying, VIC activation, and apoptosis genes using group-averaged counts per million (CPM) values that were z-scored per gene across sex *x* group combinations (**Figure 6a**). PAB C-Males showed the highest relative expression of multiple chemokines and inflammatory mediators (CXCL10, CXCL8, SPP1, CXCL14), matricellular and ECM genes (THBS4, SPARC, COL3A1, COL4A1), matrix-modifying enzymes (ADAMTS4, ADAMTS9, LOXL1, LOXL2), and apoptosis-related genes (BCL3, CFLAR, BCL2, CDKN1A), indicating a broad inflammatory and matrix-remodeling program. In contrast, PAB NC-Males most strongly expressed ECM structural and VIC activation-associated genes – including SPARC, LOXL1, LOXL2, SPP1, ACTA2, and collagen isoforms such as COL1A2, COL3A1, and COL4A1 – with comparatively less widespread induction of immune mediators. Female leaflets occupied the lower end of the expression range for all genes in this inflammatory and ECM-remodeling panel and had the smallest PAB-induced DEG set, prompting us to ask whether this modest transcriptional response was organized into a focused ECM program.

**Figure 6.**
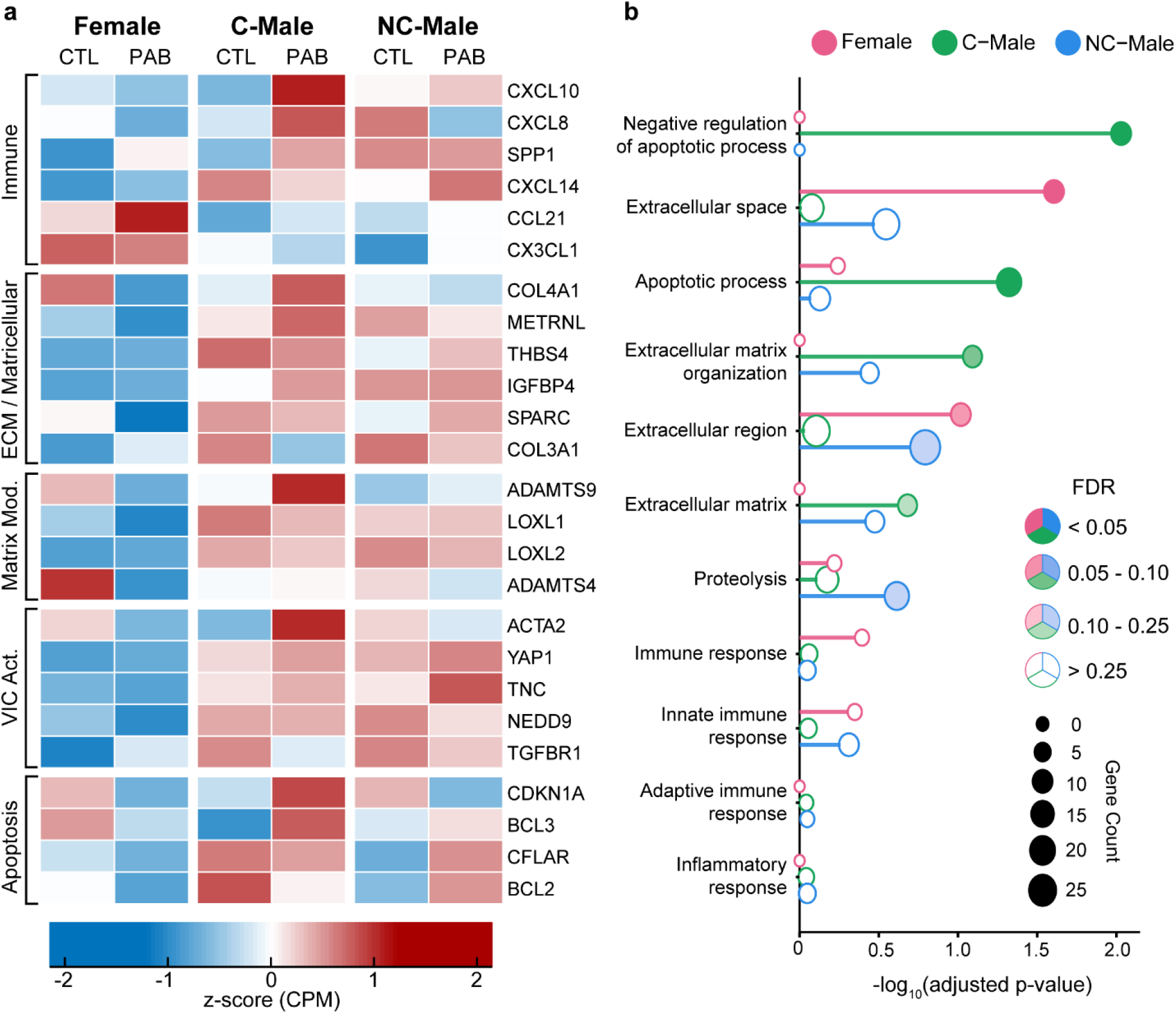
Sex–specific extracellular matrix remodeling signature and enriched biological pathways in the pressure–overloaded tricuspid valve. (a) Heatmap of log2 counts per million (CPM) for a curated panel of immune, extracellular matrix (ECM)/matricellular, matrix-modifying (Matrix Mod.), valve interstitial cell activation–associated (VIC Act.), and apoptosis genes in tricuspid valve tissue. Expression values are averaged within sex and group combinations (Female, castrated male (C-Male), and non-castrated male (NC-Male); PAB vs. CTL) and are z-scored per gene to highlight relative differences expression across the six groups. Genes were selected a priori based on established associations with ECM structure, matrix remodeling, inflammation, and cellular activation. (b) Curated Gene Ontology (GO) biological process enrichment analysis of upregulated genes within each sex-stratified comparison. The panel focuses on ECM-, proteolysis-, apoptosis-, and immune-associated pathways. Each point represents a sex-specific enrichment result for a given pathway; point size reflects gene count and the x-axis denotes - log_10_(adjusted p-value). Pathways with FDR < 0.1 are considered significantly enriched.

To determine whether these gene-level differences reflected coordinated biological programs, we performed Gene Ontology (GO) enrichment analysis on upregulated genes within each sex-stratified comparison (FDR < 0.1; **Figure 6b**). Although they showed lower expression than male groups, Females had significant enrichment in *extracellular space* (GO:0005615, FDR = 0.025, 6 genes including METRNL, SPARC, CXCL14, CST3, IGFBP4, and GLDN) and *extracellular region components* (GO:0005576, FDR = 0.096), consistent with active ECM protein secretion. C-Males demonstrated robust enrichment in apoptosis-related pathways, including *negative regulation of apoptotic process* (GO:0043066, FDR = 0.009; 6 genes including UBE2Z, HSPB1, PTK2B, NPM1, NIBAN2, and BCL3) and *apoptotic process* (GO:0006915, FDR = 0.048; 13 genes), along with significant enrichment in *extracellular matrix organization* (GO:0030198, FDR = 0.081; 5 genes including ADAMTS4, ADAMTS9, ADAMTSL2, COL4A1, and COL11A1), consistent with a coordinated matrix-remodeling program. NC-Males, despite exhibiting the largest number of DEGs, did not reach statistical significance for any of these canonical remodeling pathways at the prespecified FDR threshold. Instead, extracellular and matrix-related terms were present among the nominal pathway enrichments but lacked the pathway-level coherence seen in C-Males, indicating a more diffuse and less coordinated response in NC-Males. In contrast, downregulated genes did not show significant pathway enrichment for any sex across ECM-, proteolysis-, apoptosis-, or immune-associated pathways (all FDR > 0.1; **Figure S15**). These findings indicate sex-specific molecular mechanisms in PAB-induced tricuspid leaflet remodeling, with female leaflets prioritizing extracellular matrix protein synthesis and secretion, C-Male leaflets engaging coordinated apoptosis-regulatory and ECM-organization programs, and NC-Male leaflets mounting a broadly distributed ECM and protease response.

## 4. DISCUSSION

Heart valve leaflets are no longer seen as passive bystanders in valvular disease progression, but rather as active mediators capable of sensing and responding to biomechanical stress [20]. Studies of the mitral and aortic valves have shown that pathological mechanical loading activates VICs, promotes ECM remodeling, and drives leaflet thickening and stiffening [10,31,32]. We replicated this tissue response in the tricuspid leaflets, highlighting the heterogeneous leaflet-specific maladaptive response to secondary TR in male subjects [15,18]. Despite the known clinical sex disparities in TR prevalence, progression, and long-term outcomes, the influence of sex and sex-steroid hormones on the intrinsic tricuspid leaflet remodeling response remains largely unexplored [3–5].

Our findings demonstrate that tricuspid leaflet maladaptation is not uniform, but rather a sex- and testosterone-associated process that produces unique remodeling patterns. Under comparable hemodynamic insult and TR severity, Females, C-Males, and NC-Males developed distinct spatial, mechanical, and transcriptional remodeling phenotypes. C-Males exhibited the greatest maladaptive response, with extensive leaflet growth, thickening, increased cellularity, and low-stretch stiffening, consistent with prior male-dominant ovine secondary TR studies [14,15,18]. Females showed a more restricted leaflet- and region-specific remodeling phenotype with localized thickness changes alongside uniform low-stretch stiffening across all leaflets. NC-Males exhibited preferential septal leaflet remodeling as shown by the significant growth, thickening, stiffening, and change in cellular nuclei count, which deviated from the anterior leaflet predominance previously reported in ovine secondary TR models [18,33]. These findings identify sex and testosterone status as important biological modulators of tricuspid leaflet remodeling.

The diverse remodeling phenotypes observed across sex groups may reflect how the RV and tricuspid valve apparatus uniquely adapt to pressure overload. The known intrinsic sex differences in right-heart geometry and composition may have influenced the adaptive response of the valvular complex under pathological loading [5,34]. Females experienced changes in RV geometry, including annular and ventricular dilation, while both Male groups preserved chamber size at the cost of systolic function. This suggests Females tolerate pressure overload through geometric adaptation, while Males progress toward contractile dysfunction, consistent with sex-dependent RV remodeling reported in rat pulmonary hypertension models [35]. These different RV responses to pressure overload may create distinct mechanical loading environments across the valve apparatus which shape leaflet remodeling behavior. In the current model of PAB, only C-Males sustained elevated pulmonary pressures, had impaired RV systolic function, and preserved chamber size. This combination likely imposed a larger and more sustained mechanical burden on the tricuspid valve. These testosterone-deficient Males exhibited the most pronounced multi-leaflet remodeling response, suggesting that low circulating testosterone may drive a more fibrotic response under chronic pressure overload in Males. This pattern aligns with clinical data linking low testosterone in male patients to a greater risk of cardiovascular disease events [24,29,36]. In contrast, Females displayed more restricted leaflet remodeling, possibly reflecting a compensatory adaptive response to altered RV geometry rather than progression toward fibrosis. Collectively, these findings suggest that tricuspid leaflet remodeling exists along a spectrum between adaptation and maladaptive fibrosis, with sex-specific cardiac structure and circulating testosterone levels influencing this balance.

The molecular signatures identified in each sex group support the presence of biologically distinct remodeling programs underlying these unique phenotypes. Recent transcriptomic studies have shown that tricuspid leaflet remodeling in secondary TR involves VIC-mediated ECM turnover, inflammatory signaling, and profibrotic pathway activation [11,20]. Our findings build on this by demonstrating that remodeling programs deviate substantially according to sex and testosterone status. In Females, leaflets mounted a focused ECM secretory response, enriched for *extracellular space* and *extracellular region*, indicating a coordinated matrix remodeling rather than widespread fibrosis [9,37]. In contrast, C-Male leaflets activated a broad apoptosis regulation and ECM remodeling program marked by the induction of BCL3, CFLAR, BCL2, BNIP3, CDKN1A, and ADAMTS-family genes, along with enrichment of *negative regulation of apoptotic pathways*. This transcriptional profile is consistent with sustained fibroblast survival and matrix accumulation under chronic overload conditions [29]. These findings support recent single-cell analyses identifying fibrosis-associated VIC subpopulations enriched for ECM-remodeling programs in diseased tricuspid valves [20,38].

Females and C-Males showed different remodeling responses despite similar circulating testosterone concentrations, indicating that testosterone is not the sole driver of these differences. This suggests sex-intrinsic regulatory mechanisms, potentially including sex chromosome-dependent control of VIC activation and matrix homeostasis, may shape how tricuspid leaflets adapt to pressure overload [39]. Overall, these data highlight sex and testosterone status in Males as important biological regulators of tricuspid leaflet maladaptation.

These findings have important implications for understanding the heterogeneous clinical progression of secondary TR and for developing sex-informed therapeutic strategies. Clinical studies consistently demonstrate sex differences in TR prevalence, RV remodeling patterns, and postoperative outcomes, but the biological basis underlying these disparities remains poorly defined [3,5]. In our model, sex-dependent RV adaptation to pressure overload created distinct loading conditions on the tricuspid valve, and the leaflets developed sex- and testosterone-specific phenotypes that differed in extent, regional distribution, mechanics, and transcriptional programs of growth and fibrosis. The identification of testosterone-sensitive remodeling programs suggests that future therapies targeting VIC activation, ECM turnover, or fibrotic signaling may require consideration of biological sex and sex-steroid hormone status to optimize outcomes.

This study provides the first multiscale, sex-stratified characterization of tricuspid leaflet remodeling to secondary TR, but several limitations should be considered. Transcriptomic analyses were performed using bulk RNA sequencing of the leaflet belly-region, which cannot fully resolve regional transcriptional heterogeneity. Future spatial and single-cell approaches will be important for defining the cellular drivers of these phenotypes [20,38]. The current PAB model produces sustained TR and chronic RV remodeling, inducing pressure overload more rapidly than the gradual progression of human secondary TR. Several datasets did not detect significant sex-disease interaction effects within the mixed-effects models, likely due to both biological complexity and limited statistical power. Additionally, testosterone was not directly manipulated in this study, and differences between C-Males and NC-Males may also be attributed to any of the downstream consequences of castration. These findings should be interpreted as evidence for sex-associated remodeling phenotypes rather than definitive categorical sex differences.

In conclusion, this study demonstrates tricuspid leaflet maladaptation during secondary TR as a sex-specific and testosterone-sensitive process spanning structural, mechanical, and transcriptional scales. By integrating multi-leaflet structural characterization with biomechanical and transcriptomic analyses, these findings advance the emerging view of the tricuspid valve as an active biological participant in disease progression. Future studies defining the mechanobiological and cellular mechanisms underlying these sex-dependent remodeling programs may help establish new approaches for predicting disease progression and developing targeted therapies for secondary TR.

## ACKNOWLEDGEMENTS

The opinions, findings, and conclusions, or recommendations expressed are those of the authors and do not necessarily reflect the views of the American Heart Association or National Institute of Health. We acknowledge the University of Texas at Austin Center for Biomedical Research Support (CBRS) Genomic Sequencing and Analysis Facility (GSAF) and Bioinformatics Team for their assistance with Tag sequencing, read deduplication, and adapter trimming. They thank Dr. Michelle Harrison and Dr. Stephanie Seidlits at The University of Texas at Austin for use of instrumentation needed for blood plasma multiplexing for this study.

## DATA AVAILABILITY

The trimmed fastq RNA sequencing files are available in the National Center for Biotechnology Information Sequence Read Archive under BioProject PRJNA1505704. All other data will be uploaded to the Texas Data Repository and become available upon publication.

## AUTHOR CONTRIBUTIONS

**Colton J. Kostelnik:** Conceptualization, Data curation, Software, Formal analysis, Investigation, Methodology, Writing -- original draft, Writing -- review & editing. **Magda L Piekarska:** Conceptualization, Data curation, Investigation, Methodology, Writing -- review & editing. **Shreya Sreedhar:** Data curation, Investigation, Writing -- review & editing. **Chien-Yu Lin:** Investigation, Writing -- review & editing. **Aarya Shah:** Data curation, Investigation, Formal analysis. **Boguslaw Gaweda:** Investigation, Methodology, Writing -- review & editing. **Austin Goodyke:** Writing - review & editing. **Xu Yujun:** Software, Validation, Writing --review & editing. **Kartik Balachandran:** Writing – review & editing. **Layla Parast:** Validation, Writing - review & editing. **Matthew R. Bersi:** Software, Validation, Writing -- review & editing. **Tomasz A. Timek:** Conceptualization, Resources, Supervision, Funding acquisition, Writing -- review & editing. **Manuel K. Rausch:** Conceptualization, Writing -- original draft, Writing -- review & editing, Supervision, Funding acquisition.

