## Supplemental Materials for "Sex–Specific Remodeling Phenotypes of the Tricuspid Valve Leaflets in an Ovine Model of Functional Tricuspid Regurgitation"

**Supplemental Methods:**

***Animal Model, Medication, and Procedures***

A All experiments complied with the Principles of Laboratory Animal Care and the National Institutes of Health Guide for the Care and Use of Laboratory Animals and were approved by the Michigan State University Institutional Animal Care and Use Committee (protocols PROTO202200120 approved 5/16/2022; and PROTO202500129 approved 5/14/2025).

Healthy adult Dorset sheep were randomly assigned to either control (CTL) or pulmonary artery banding (PAB) treatment groups, with all male sheep obtained as castrated and non-castrated from Michigan State University. Final group sample sizes are shown in **Table 1**. Animals were anesthetized with guaifenesin (50-100ml of 5% solution IV), lidocaine (1mg/kg IV), and propofol (4-6 mg/kg IV). In PAB animals, we then performed a sterile limited left thoracotomy through the fourth intercostal space and recorded pulmonary artery pressures using direct measurement with a 20 G catheter inserted in the proximal pulmonary trunk. Epicardial echocardiography (Vivid S6, GE Healthcare; 1.5–3.6 MHz transducer) was used to assess ventricular function and valvular competence at baseline and terminal studies. We evaluated valvular insufficiency based on the American Society of Echocardiography guideline, with TR severity graded by a specialized veterinary echocardiographer from 0 (none) to 4 (severe) based on color Doppler jet area and vena contracta width. To induce RV dilatation and remodeling, the pulmonary artery was encircled with an umbilical tape band at the main distal pulmonary trunk, progressively tightened to achieve a target systolic pulmonary arterial pressure at least double that of baseline, and then secured in place with surgical clips [1,2]. For postoperative infection prophylaxis, all animals received cefazolin (2 g

IV or IM every 12 hours) and gentamicin (240 mg IV every 24 hours) for 10 days postoperatively, beginning with a preoperative dose administered prior to incision. Animals were then recovered and followed for 12–15 weeks after PAB before terminal studies (Supplemental Table S1); CTL animals did not undergo PAB or sham thoracotomy.

Following the recovery period, we performed terminal hemodynamic and echocardiographic measurements on all secondary TR induced sheep in the same manner as the initial PAB surgery but through a median sternotomy to optimize access and echocardiographic windows. We then euthanized all animals with sodium pentothal (100 mg/kg IV) and potassium chloride (80 mEq IV). For CTL sheep, we collected terminal epicardial echocardiography and hemodynamic measurements through a median sternotomy before euthanizing the animals. After the animals were euthanized, we removed the heart and isolated the tricuspid valve complex from each animal for further analysis.

***Plasma Multiplexing***

We collected whole blood from each animal into EDTA-treated tubes and centrifuged at 10,000 x g for 10 minutes. We aliquoted the plasma and stored the samples at -80°C until analysis. Plasma cytokine (IL-1$\beta$, IL-6, IL-8, IL-10, and TNF-$\alpha$) concentrations were measured using the Ovine Cyto/Chemokine Panel 1 (MilliporeSigma, Burlington, MA, USA) and plasma hormone (testosterone, estradiol, progesterone, cortisol) concentrations were measured using the Multi-Species Mag Panel (MilliporeSigma, Burlington, MA, USA), following the manufacturers' instructions. We measured each plate on a Luminex xMAP-Intelliflex system and fit the standards using the Belysa Immunoassay Curve Fitting Software.

***Leaflet Morphology and Storage***

We carefully removed the tricuspid valve complex from each explanted ovine heart to avoid leaflet damage by first making a 1 cm atrial incision parallel to the atrioventricular groove. We then rinsed all heart chambers with 1x PBS to remove any residual blood and prevent coagulation on the leaflets.

Next, we opened the right ventricle (RV) by cutting from the posterior–septal commissure toward the apex parallel to the interventricular septal plane and cut the chordae tendineae at their papillary muscle insertions to prevent tension on the leaflets. Lastly, we cut circumferentially along the annulus to remove the tricuspid leaflets.

We then floated the intact valves in 1x PBS and photographed each valve on a calibrated grid to measure and compare morphological parameters, including leaflet area, between CTL and PAB animals (**Figure 1a**). Following imaging, we separated the valve into the three major leaflets and cryopreserved each sample in DMEM with 10% DMSO supplemented with a protease inhibitor (ThermoFisher, A32953, Waltham, MA, USA). We froze the samples at a controlled rate of 1 °C/min and stored them at −80 °C until further testing to minimize changes in cellular and mechanical properties [3,4].

***Leaflet Profilometry***

A fter photographing the thawed leaflets, we gently dried each using absorbent tissue. Next, we covered the leaflets with microfine talc granules (ThermoFisher, A0448752, Weltham, MA, USA) to minimize glare and reduce the tissue's optical transparency. Lastly, we imaged the leaflets using an optical profilometer (VHX5000, Keyence, Osaka, Japan) under 50× magnification (**Figure 2a**). Immediately following the 3D scanning, we washed the tissue in 1xPBS supplemented with 1 U/mL RNase inhibitor (SUPERase-In, Invitrogen, Thermo Fisher Scientific, Waltham, MA, USA) to preserve RNA integrity. We have shown that this does not negatively impact the biomechanical characterization of leaflets [5]. To compare whole-leaflet thickness changes across treatment groups and sexes, we averaged the thickness across the entire leaflet surface and report absolute thickness values. To assess regional thickness changes between treatment groups, we meshed each thickness map, transformed each map into a fine matrix preserving native spatial resolution, then averaged values into a coarse 3 x 3 matrix, with rows representing radial position (near annulus, belly, free edge) and columns representing circumferential position spanning the leaflet's two commissures (**Supplemental Figure S6**). This approach allowed us to group regional thicknesses across leaflets of varying geometries while preserving anatomical location. We then performed a log2-fold change (FC) comparison of the regional thicknesses between treatment groups stratified by sex and leaflet.

***Leaflet Biaxial testing***

We prepared a mechanical testing sample from each septal, anterior, and posterior leaflet by excising a 7 x 7 mm square sample from the central belly region of each leaflet (**Figure 3a**). We then placed four fiducial markers on the atrial surface of each sample to track local strain throughout testing. Prior to mounting, we measured the sample thickness at four sites with a digital thickness gauge (547-500S, Mitutoyo Corp.), and we photographed each specimen in a stress-free configuration on a calibrated grid.

We mounted samples on a commercial biaxial testing device (Biotester 5000, CellScale Biomaterials Testing, Waterloo, ON, Canada) and submerged them in 1xPBS at 37°C. The mechanical test began with ten equibiaxial preconditioning cycles to 300 mN to stabilize the mechanical response, consistent with our earlier ovine tricuspid leaflet studies [6,7]. After preconditioning, we applied two force-controlled equibiaxial cycles to 300 mN at an approximate loading rate of 23 mN/s. We limited the protocol to 300 mN to capture the full nonlinear behavior without damaging the tissue, which we observed near 400 mN. Throughout loading, we continuously recorded circumferential and radial forces as well as images of the fiducial markers at a rate of 5 Hz.

We analyzed the images using the software LabJoy (CellScale Biomaterials Testing, Waterloo, ON, Canada) to track coordinates of the fiducial markers throughout testing. We analyzed the unloading curve of the final equibiaxial cycle, and calculated membrane tension-stretch data relative to the stress-free configuration. This stress-free configuration provided a consistent reference state and avoided artificial stretch introduced during mounting [8]. We tracked four fiducial markers on each leaflet between the stress-free reference configuration and each deformed (loaded) configuration. We computed the deformation gradient tensor, $\boldsymbol{F}$, from the marker displacements relative to the stress-free reference state, and then derived the right Cauchy-Green deformation tensor $\boldsymbol{C} = \boldsymbol{F}^T\boldsymbol{F}$. We obtained circumferential and radial stretches as the square roots of the corresponding diagonal components, $\boldsymbol{\lambda_{circumferential}} = \sqrt{\boldsymbol{C_{11}}}$ and $\boldsymbol{\lambda_{radial}} = \sqrt{\boldsymbol{C_{22}}}$. We then computed membrane tension by dividing the measured force by the perpendicular rake-to-rake distance in the deformed configuration at each time point.

To compare the nonlinear membrane tension–stretch curves between groups, we identified three characteristic parameters (**Supplemental Figure 8b**): (i) toe stiffness, (ii) calf stiffness, and (iii) the transition stretch. Toe stiffness was defined as the tangent modulus of the low-tension region of the curve, identified as the largest contiguous segment starting from the origin for which a linear fit achieved $R^2 \geq 0.99$. Calf stiffness was defined as the tangent modulus at the membrane tension nearest to 20 N/m, within the physiological loading range of the tricuspid valve leaflets. The transition stretch was defined as the stretch ratio at the intersection of the toe and calf stiffness linear fits, identified as the closest sampled data point to that intersection.

***Leaflet Histology***

We fixed full-length radial strips (from the annulus to the free edge) of all leaflets in 10% neutral buffered formalin for 2 hours prior to transferring the tissue to 70% ethanol for long-term storage. We then shipped the fixed leaflet strips to a commercial histology service (HistoServ Inc., Amaranth, MD) for paraffin embedding, sectioning at 5 µm, and Hematoxylin & Eosin (H&E) staining. We acquired high-content full section images for all H&E-stained leaflets using a Nikon Eclipse Ti2-E inverted microscope equipped with NIS-Elements HC software (Nikon Corporation, Tokyo, Japan) at 10x magnification. To focus our analysis on the leaflet, we manually excluded annular muscle bundles by masking regions containing striated muscle based on H&E morphology. To quantify regional cellular density, we fit splines to each leaflet's atrial and ventricular surfaces to define the tissue boundary, then used these boundaries to segment the cross-sectional image into three equidistant radial regions (near annulus, belly, free edge) and three transmural bands (atrial, mid, ventricular) to balance spatial resolution with statistical power (**Supplemental Figure S11a**). We used hue, saturation, and value (HSV) thresholding to identify cell nuclei stained with hematoxylin (H: 218-337, S: 0-1, V: 0-0.78), building on prior work using HSV color space for histologic nuclei segmentation [6,7]. Using these thresholds, we quantified the total nuclei count for the full leaflet cross section and compared absolute counts between control and diseased groups for each sex and leaflet. We also calculated regional cell density as the number of nuclei per unit area within each region and report values as the log2(FC) in cell density between respective control and diseased groups for each region, leaflet, and sex.

***Leaflet RNA Sequencing***

We excised a 6 mm biopsy punch from the leaflet belly regions immediately following 3D profilometry. The biopsy samples were preserved in RNAlater Stabilization Solution (Invitrogen, Thermo Fisher Scientific, Waltham MA, USA) prior to total RNA extraction using the RNeasy Fibrous Tissue Mini Kit (QIAGEN, Hilden, Germany). RNA integrity was assessed in a representative subset of samples with a median RNA integrity number (RIN) of 7.6, indicating adequate preservation of RNA for bulk RNA sequencing [9]. We then submitted the total RNA to The University of Texas at Austin Genomic Sequencing and Analysis Facility for library preparation and 3' Tag Sequencing (TagSeq) [10]. Libraries were prepared, quality-controlled, and pooled according to standard TagSeq protocols before sequencing on a NovaSeq 6000 (RRID:SCR_016387) (Illumina, San Diego, CA, USA) with single-end 100-bp reads.

TagSeq library deduplication and adapter trimming were performed by The University of Texas at Austin Computational Biology and Bioinformatics Core Facility (RRID: SCR_022688). All computational analyses were performed on the Texas Advanced Computing Center (TACC) high-performance cluster. We aligned high-quality reads to the Ovis aries reference genome ARS-UI_Ramb_v3.0 (RefSeq assembly GCF_016772045.2; NCBI) using STAR (v2.7.11b). We assigned aligned reads to annotated gene features using HTSeq-count (v2.0.7), and then performed differential gene expression using DESeq2, with group (CTL vs. PAB) and sex included as fixed effects with interactions. We normalized gene counts using regularized log transformation and applied a Benjamini–Hochberg correction, considering genes with an adjusted $p \leq 0.05$ and $|\log2(FC)| \geq 0.5$ as significant. Gene Ontology (GO) biological process enrichment analysis was performed using clusterProfiler (version 4.18.4) with Benjamini-Hochberg false discovery rate (FDR) correction; significantly upregulated genes (adjusted $p \leq 0.05$ and $\log2(FC) > 0.5$) from each sex-stratified comparison were used as input gene lists. Pathways with FDR $\leq 0.1$ were considered significantly enriched.

***Statistical Analysis***

Forty-five sheep were used in this study and were age-matched across sexes within each group. Body weights differed by sex (**Supplemental Table S1**). To account for this, we indexed echocardiographic right heart by body weight, and included body weight as a fixed-effect covariate in our statistical models. We excluded body weight from regional cell density, since we normalized nuclei count by regional cross-sectional area and did not expect it to scale with body size. We analyzed morphological, thickness, biaxial, and histological data using linear mixed-effects models in R (*lme4* package). Common fixed effects across models included group (CTL vs. PAB), sex (Female, C-Male, NC-Male), and leaflet (anterior, posterior, septal), along with their interactions to assess sex-dependent and leaflet-specific remodeling. We incorporated additional fixed effects as appropriate for specific datasets: for biaxial data, we included tissue direction (circumferential vs. radial) along with its interactions with group, sex, leaflet to evaluate direction-dependent remodeling, while regional thickness and nuclei density models included spatial location (radial and circumferential regions) to enable region-specific comparisons. All models included a random effect for subject to account for repeated measurements within animals, and we used Type III ANOVA with Satterthwaite's approximation to assess the significance of fixed effects and interactions.

We conducted post-hoc pairwise comparisons using estimated marginal means (*emmeans* package). We evaluated group differences (CTL vs. PAB) within each leaflet and sex (and tissue direction, where applicable), and compared sexes within each group-leaflet combination. For regional data, we performed post-hoc pairwise comparisons within each combination of leaflet, sex, and spatial region, adjusting p-values with the FDR correction. We defined statistical significance as $p < 0.05$, and we report data as mean ± standard deviation.

**<u>Supplementary Tables</u>:**

**Table S1.** Terminal body weight, age, and banding duration by sex and disease status.

| | **Female** | | **Castrated Male (C-Male)** | | **Non-castrated Male (NC-Male)** | |
|---|---|---|---|---|---|---|
| | **CTL (n = 9)** | **PAB (n = 10)** | **CTL (n = 7)** | **PAB (n = 9)** | **CTL (n = 5)** | **PAB (n = 5)** |
| Weight (kg) | 54.5 ± 4.1[#,§] | 65.7 ± 4.1[*,#,§] | 68.4 ± 4.2 | 71.6 ± 6.3 | 69.8 ± 3.6 | 75.8 ± 3.8 |
| Age (months) | 8.0 (4.0) | 8.0 (1.0) | 8.3 ± 1.4 | 10.0 ± 1.9[*] | 9.0 (0.0) | 9.0 (0.0) |
| Weeks Banded | - | 12.9 (2.7) | - | 12.4 (3.3) | - | 12.0 (3.1) |

Values are mean ± standard deviation or median (interquartile range) as determined by the Shapiro-Wilk normality test. Statistical differences were determined from pairwise comparisons of linear mixed effect model.

CTL = control, PAB = pulmonary artery banding.

* $p < 0.05$ (CTL vs. PAB), # $p < 0.05$ (Female vs. C-Male), ¥ $p < 0.05$ (C-Male vs. NC-Male), § $p < 0.05$ (Female vs. NC-Male)

**Table S2.** Hemodynamic measurements for pulmonary artery banded (PAB) castrated male (C-Male), non-castrated male (NC-Male), and female sheep at three timepoints: baseline, acutely following the PAB procedure (Post-PAB), and prior to euthanasia (Terminal).

| | **Female** | | | **Castrated Male (C-Male)** | | | **Non-castrated Male (NC-Male)** | | |
|---|---|---|---|---|---|---|---|---|---|
| | **Baseline (n = 10)** | **Post-PAB (n = 10)** | **Terminal (n = 10)** | **Baseline (n = 9)** | **Post-PAB (n = 9)** | **Terminal (n = 8)** | **Baseline (n = 5)** | **Post-PAB (n = 5)** | **Terminal (n = 5)** |
| sPAP, mmHg | 14 (11.5) | 54.7 ± 3.6[‡,†] | 27.1 ± 4.7 | 15.8 ± 3.6 | 52.6 ± 6.2[‡,†] | 31.0 (17.8)[€,#] | 17.6 ± 4.8 | 58.8 ± 3.3[‡,†] | 36.6 ± 7.5 |
| dPAP, mmHg | 8.9 ± 4.2 | 13.0 (7.0) | 9.0 ± 4.3 | 6 (3) | 11.4 ± 5.4 | 11.5 (18.3)[¥] | 8.8 ± 3.0 | 11 (3.0) | 6.2 ± 0.84 |
| mPAP, mmHg | 11.8 ± 5.0 | 31.1 ± 2.9[‡,†] | 17.1 ± 3.6[#] | 11 ± 3.5 | 29.1 ± 4.8[‡] | 20.9 (16.4)[€,¥] | 12.8 ± 4.4 | 34.8 ± 4.9[‡,†] | 19.8 ± 2.3 |
| sAP, mmHg | 94.3 ± 13.2 | 101.3 ± 13.7 | 88.2 ± 12.8 | 101.1 ± 10.8 | 92.3 ± 8.9 | 77.9 ± 25.6[¥] | 97.2 ± 7.9 | 94.6 ± 6.5 | 98.2 ± 19.7 |
| dAP, mmHg | 60.2 ± 9.9 | 67.5 ± 9.0 | 67.1 ± 12.5 | 76.3 ± 18.6 | 67.8 ± 14.3 | 57.8 ± 30.0 | 69.0 (5.0) | 61.2 ± 10.2 | 77.8 ± 16.6 |
| mAP, mmHg | 72.6 ± 9.6 | 78.8 ± 10.3 | 76.5 ± 12.5 | 85.4 ± 15.4 | 77.0 ± 12.7 | 66.4 ± 27.0[¥] | 78.0 (3.0) | 72.4 ± 8.5 | 86.6 ± 16.8 |
| TR (0-4) | 0.0 (1.0) | 1.25 (1.0)[‡,†] | 4.0 (1.75)[€] | 0.0 (1.0) | 2.0 (0.0)[‡,†] | 3.5 (1.0)[€] | 0.0 (1.0) | 2.2 ± 1.1[‡] | 2.6 ± 1.3[€] |

Values are mean ± standard deviation or median (interquartile range) as determined by the Shapiro-Wilk normality test. Statistical differences were determined from pairwise comparisons of linear mixed effect model.

CTL = control, PAB = pulmonary artery banding, PAP = pulmonary artery pressure, AP = arterial pressure, s = systolic, d = diastolic, m = mean, TR = tricuspid regurgitation

**‡** $p < 0.05$ (Baseline vs. Post-PAB), **†** $p < 0.05$ (Post-PAB vs. Terminal), **€** $p < 0.05$ (Baseline vs. Terminal),
**#** $p < 0.05$ (Female vs. C-Male), **¥** $p <0.05$ (C-Male vs. NC-Male), **§** $p<0.05$ (Female vs. NC-Male)

**Supplementary Figures**:

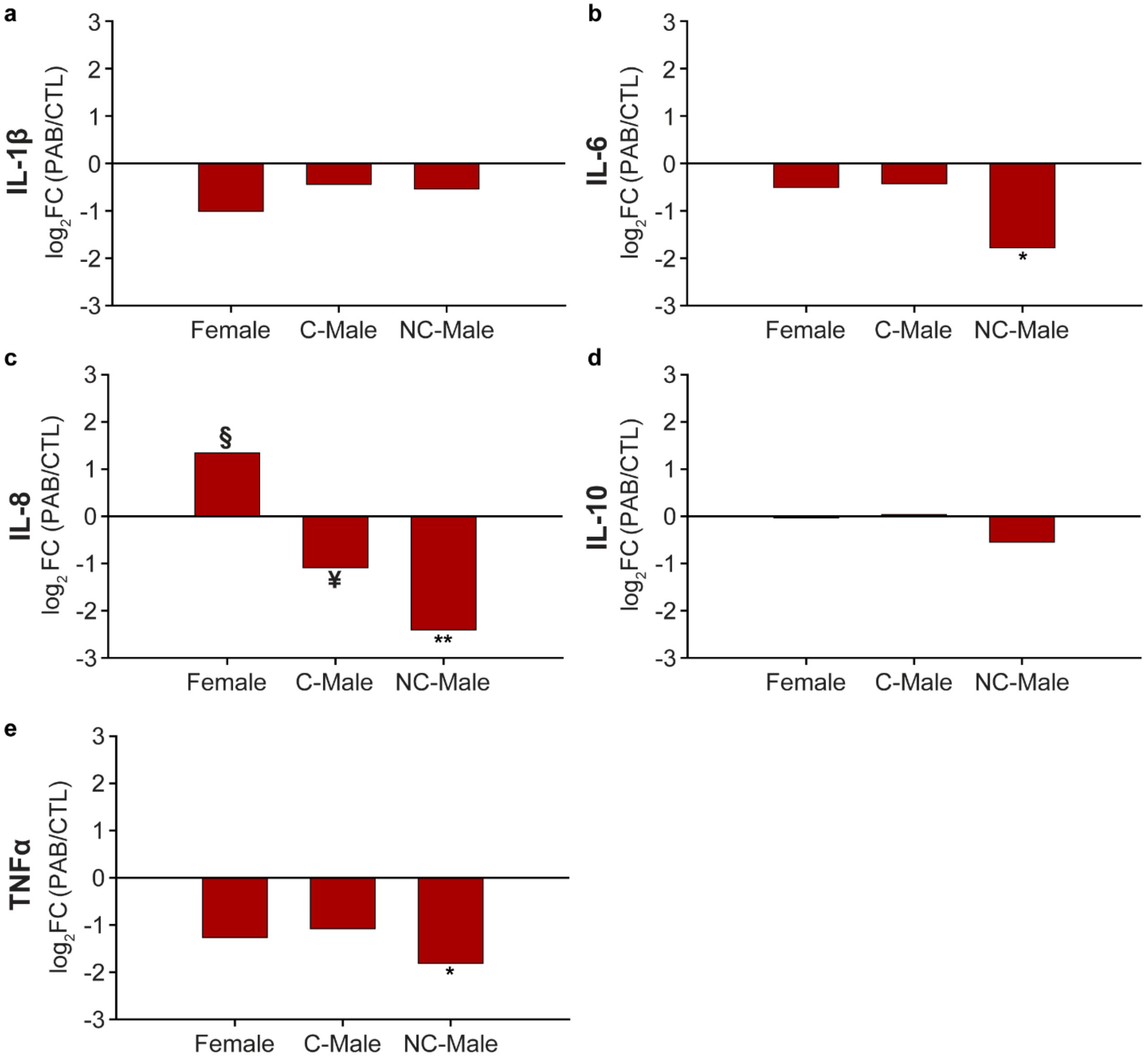


**Figure S1:** Terminal plasma cytokine log2 fold-change in pulmonary artery banding (PAB) versus control (CTL) sheep. Log2 fold-change (PAB/CTL) for (a) IL-1β, (b) IL-6, (c) IL-8, (d) IL-10, and (e) TNFα in females, castrated males (C-Male), and non-castrated males (NC-Male). Fold-change values are reported as log2(PAB/CTL) based on terminal plasma cytokine concentrations within each sex group. A value of 0 indicates no change; positive values indicate higher cytokine levels in PAB relative to CTL, and negative values indicate lower levels. Statistical comparisons between PAB and CTL within each sex group were evaluated using linear mixed-effects models with post hoc pairwise comparisons. Statistically significant contrasts between animal groups (CTL vs. PAB) are denoted by (*) for $p < 0.05$ and (**) for $p < 0.01$, and significant contrasts between sex groups are denoted by (§) for $p < 0.05$ (Female vs. NC-Male) and (¥) for $p < 0.05$ (C-Male vs. NC-Male).

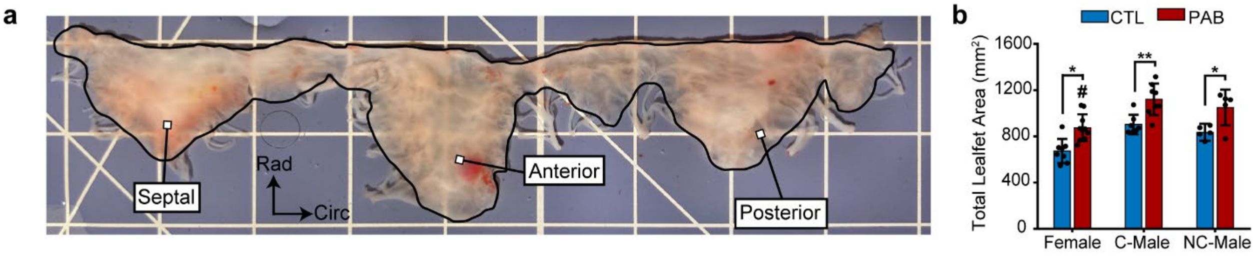


**Figure S2.** Total tricuspid valve leaflet area increases after pulmonary artery banding (PAB) for all sex groups. (a) Representative tricuspid valve leaflet on a calibrated grid (1-cm scale) with the measured area (black line) and the three main leaflets (septal, anterior, and posterior) are shown. (b) Total leaflet area comparison between control (CTL) and PAB stratified by sex (female, castrated male (C-Male), and non-castrated male (NC-Male)) sheep. Total leaflet area was calculated as the sum of individual leaflets for each animal, and data are shown for individual animals with group means ± standard deviation. Statistical differences between groups were assessed using linear mixed-effects models with post hoc pairwise comparisons of estimated marginal means. Significant contrasts between animal groups (CTL vs. PAB) are denoted by (*) for $p < 0.05$ and (**) for $p < 0.01$, and significant contrasts between sex groups are denoted by (#) for $p < 0.05$ (C-Male vs. Female).

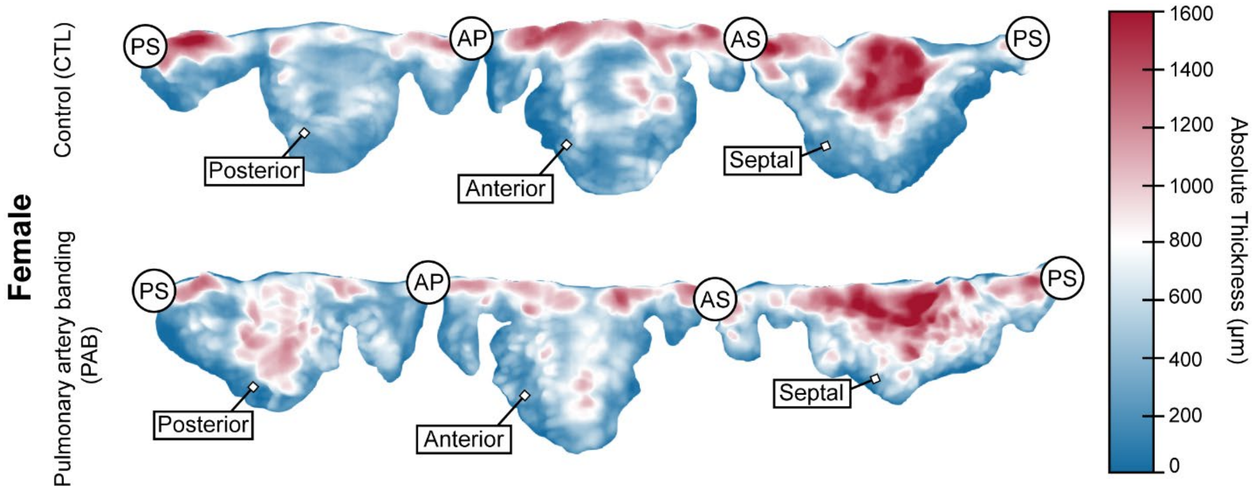


**Figure S3**. Representative thickness maps of the tricuspid valve ventricular surface in Female control (CTL) and pulmonary artery banding (PAB) animals. The posterior, anterior, and septal leaflets are labeled, along with the three commissures separating them: antero-posterior (AP), antero-septal (AS), and posterior-septal (PS). Color indicates absolute leaflet thickness in micrometers.

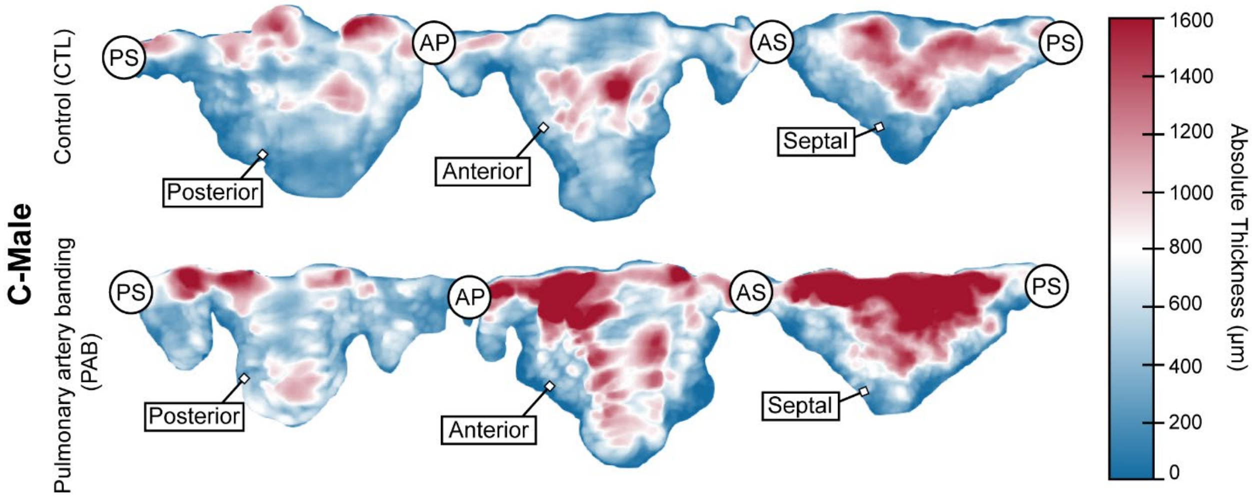


**Figure S4**. Representative thickness maps of the tricuspid valve ventricular surface in castrated male (C-Male) control (CTL) and pulmonary artery banding (PAB) animals. The posterior, anterior, and septal leaflets are labeled, along with the three commissures separating them: antero-posterior (AP), antero-septal (AS), and posterior-septal (PS). Color indicates absolute leaflet thickness in micrometers.

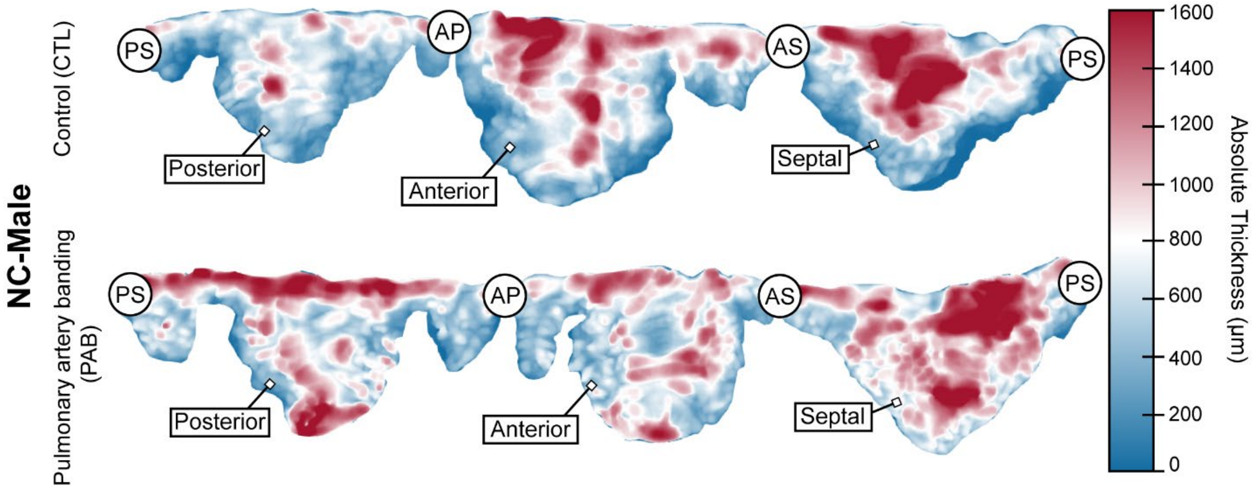


**Figure S5**. Representative thickness maps of the tricuspid valve ventricular surface in non-castrated male (NC-Male) control (CTL) and pulmonary artery banding (PAB) animals. The posterior, anterior, and septal leaflets are labeled, along with the three commissures separating them: antero-posterior (AP), antero-septal (AS), and posterior-septal (PS). Color indicates absolute leaflet thickness in micrometers.

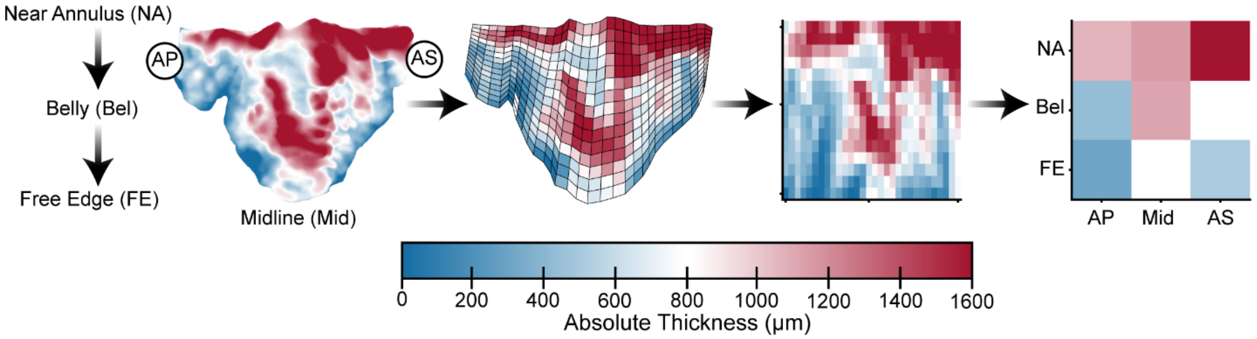


**Figure S6:** Workflow for converting a leaflet thickness map into a coarse regional matrix. Representative thickness map of a female anterior leaflet's ventricular surface after pulmonary artery banding, with regions labeled by radial zone (near-annulus (NA), belly (Bel), free-edge (FE)) and circumferential zone (antero-posterior (AP) commissural region, midline (Mid), and antero-septal (AS) commissural region). The profilometric map was discretized onto a 2D mesh, with each element retaining average 3D thickness information, then transformed into a fine matrix (rows: radial position, NA to FE; columns: circumferential position, AP to AS), which was averaged into a coarse 3x3 matrix for regional comparisons.

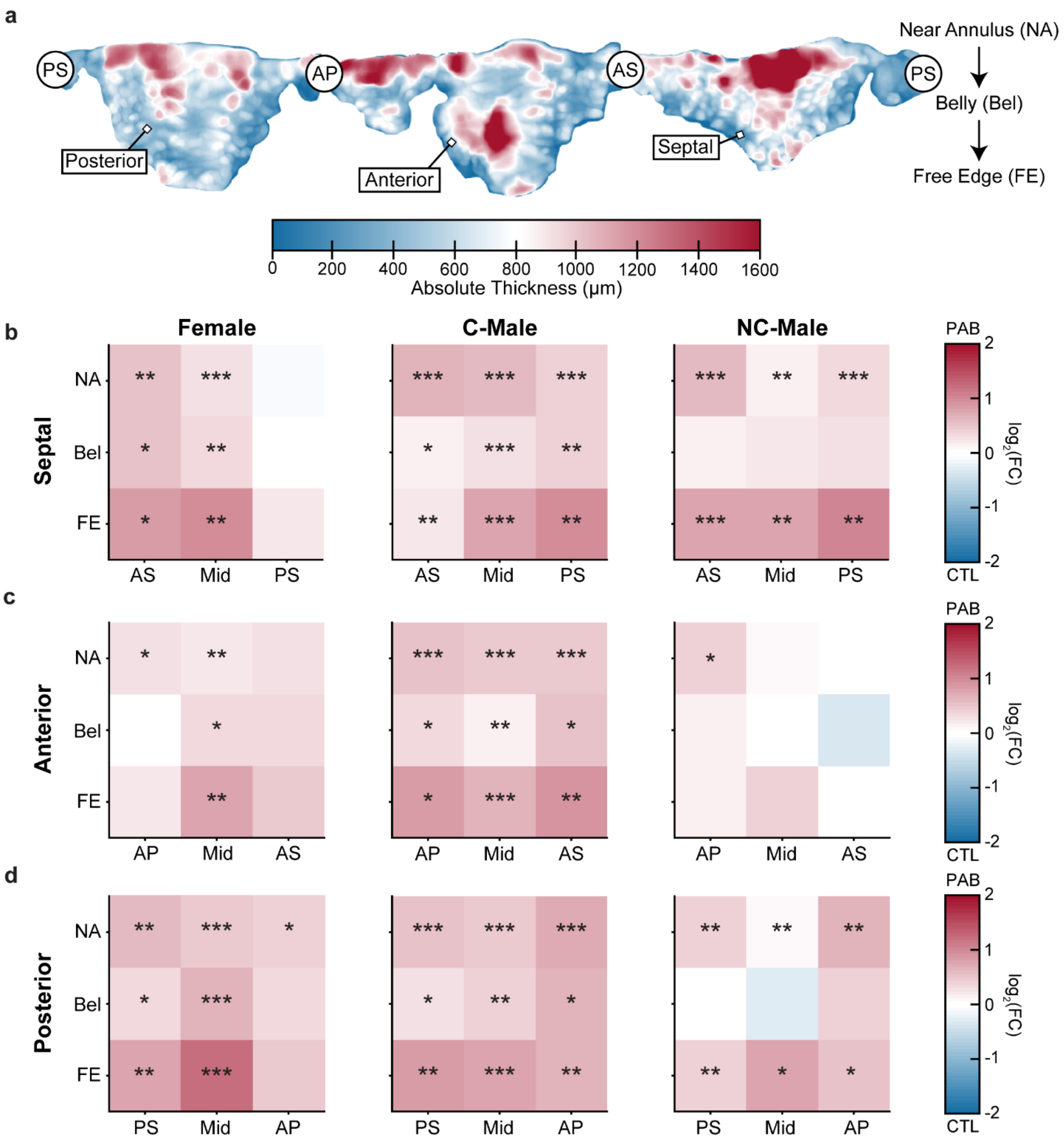


**Figure S7:** Regional tricuspid leaflet thickening after pulmonary artery banding (PAB) is greatest near the free edge and varies by sex and leaflet. (a) Representative three-dimensional thickness maps of PAB tricuspid valve leaflets with commissures labeled as posterior-septal (PS), antero-septal (AS), and antero-posterior (AP). Regional thickness fold change (FC) between PAB and control (CTL) was quantified from profilometric maps for the (b) septal, (c) anterior, and (d) posterior leaflets of female, castrated male (C-Male), and non-castrated male (NC-Male) sheep. Each leaflet profilometric map was meshed, divided radially into near-annulus (NA), belly (Bel), and free-edge (FE) regions, and divided circumferentially into three segments that correspond to each leaflet's respective commissures. The heat-map color scale reports log2(FC), interpreted as (positive, red): PAB is thicker than CTL, (0, white): PAB and CTL thicknesses are approximately equal, and (negative, blue): PAB thickness is less than CTL. Regional statistical differences between groups were assessed using linear mixed-effects models with post hoc pairwise comparisons of estimated marginal means. Significant contrasts between animal groups (CTL vs. PAB) are denoted by (*) for $p < 0.05$, (**) for $p < 0.01$, and (***) for $p < 0.001$.

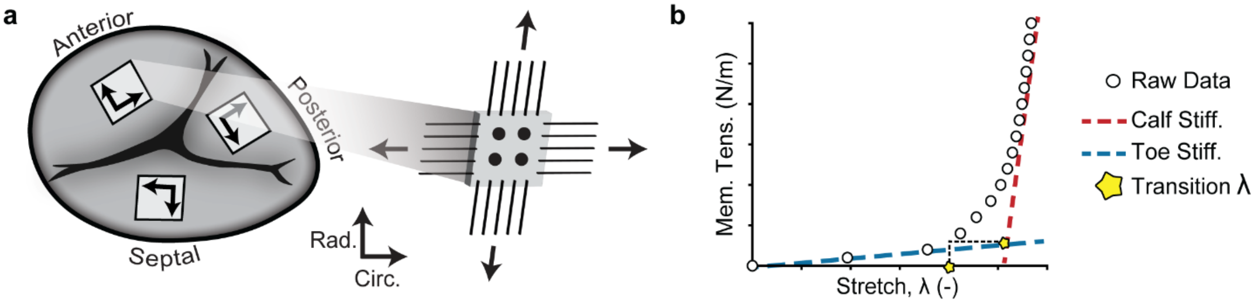


**Figure S8.** (a) Schematic of a tricuspid valve leaflet showing square regions in the anterior, septal, and posterior leaflets from which biaxial specimens were excised from the leaflet belly for testing in the radial (Rad.) and circumferential (Circ.) directions. (b) Representative membrane tension–stretch curve illustrating the mechanical metrics: open circles denote the raw data, the toe stiffness (Toe Stiff.) is defined as the slope at low stretches, the calf stiffness (Calf Stiff.) as the slope at high stretches, and the transition stretch (λ) as the stretch at which the curve transitions between these regions (heel of the J-shaped curve).

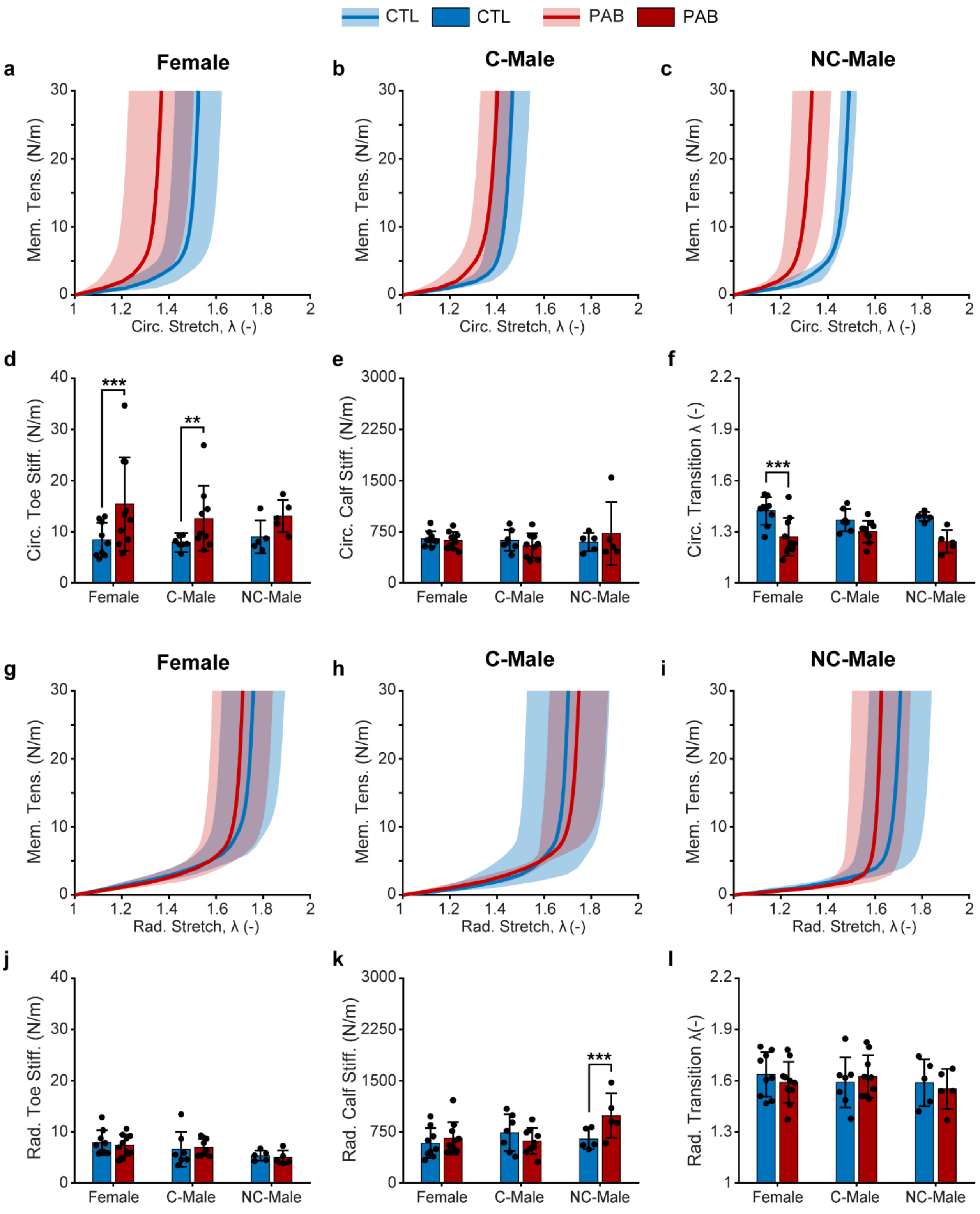


**Figure S9.** Pulmonary artery banding (PAB) alters the biaxial mechanical behavior of the septal leaflet. Circumferential membrane tension (Mem. Tens.) and stretch curves for control (CTL, blue) and PAB (red) sheep are shown for (a) females, (b) castrated males (C-Male), and (c) non-castrated males (NC-Male), with shaded regions denoting mean (solid line) and standard deviation (shaded). Corresponding circumferential mechanical metrics are presented as (d) toe stiffness, (e) calf stiffness, and (f) transition stretch (λ) for each sex group. Radial membrane tension–stretch curves for CTL and PAB sheep are shown for (g) females, (h) C-Males, and (i) NC-Males, with shaded regions denoting ± standard deviation, and the associated radial (j) toe stiffness, (k) calf stiffness, and (l) transition stretch (λ). Data are shown for individual animals with group means ± standard deviation. Statistical differences between groups were assessed using linear mixed-effects models with post hoc pairwise comparisons of estimated marginal means. Significant contrasts between animal groups (CTL vs. PAB) denoted by (*) for $p < 0.05$ and (**) for $p < 0.01$.

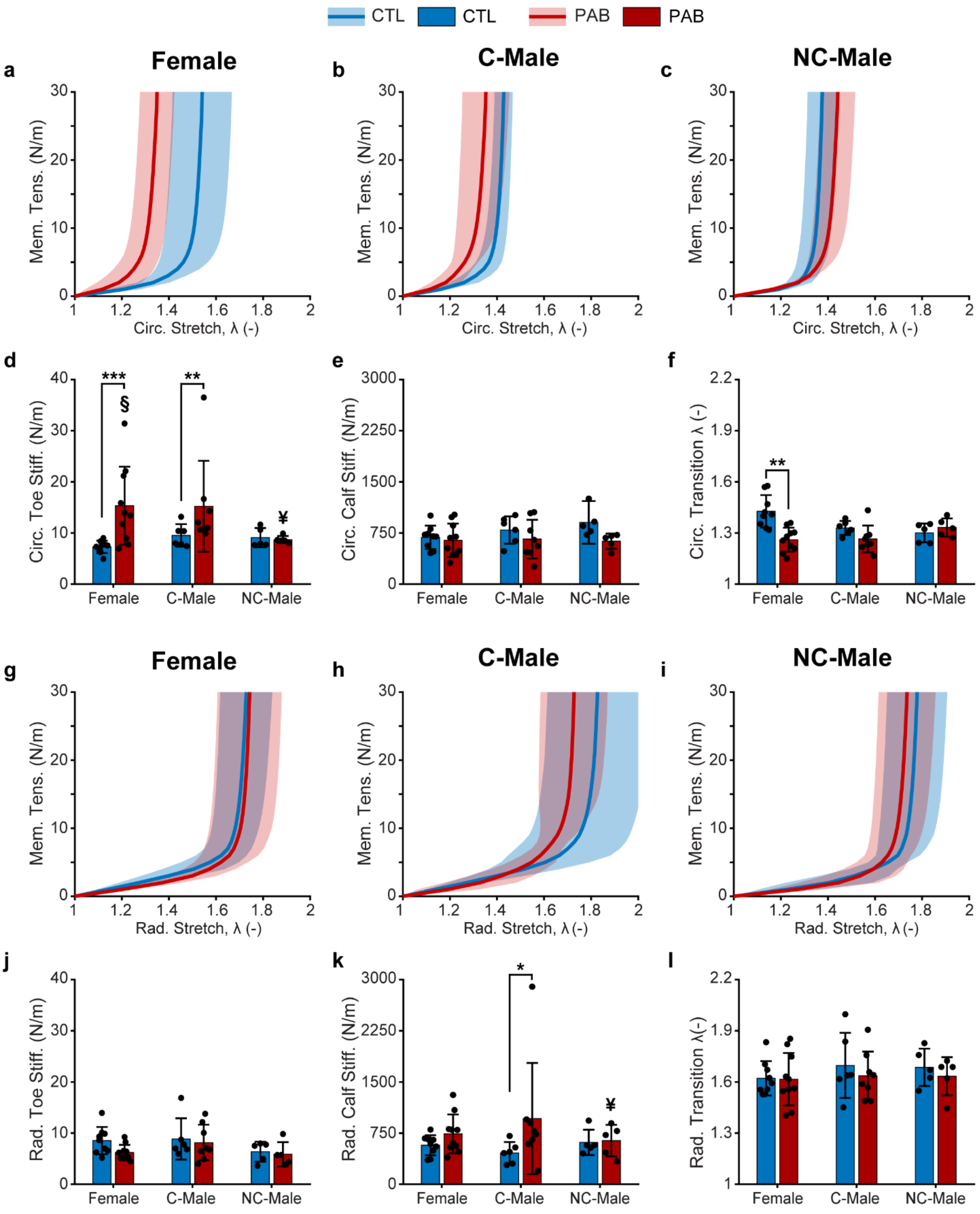


**Figure S10.** Pulmonary artery banding (PAB) alters the biaxial mechanical behavior of the posterior leaflet. Circumferential membrane tension (Mem. Tens.) and stretch curves for control (CTL, blue) and PAB (red) sheep are shown for (a) females, (b) castrated males (C-Male), and (c) non-castrated males (NC-Male), with shaded regions denoting mean (solid line) and standard deviation (shaded). Corresponding circumferential mechanical metrics are presented as (d) toe stiffness, (e) calf stiffness, and (f) transition stretch (λ) for each sex group. Radial membrane tension–stretch curves for CTL and PAB sheep are shown for (g) females, (h) C-Males, and (i) NC-Males, with shaded regions denoting ± standard deviation, and the associated radial (j) toe stiffness, (k) calf stiffness, and (l) transition stretch (λ). Data are shown for individual animals with group means ± standard deviation. Statistical differences between groups were assessed using linear mixed-effects models with post hoc pairwise comparisons of estimated marginal means. Significant contrasts between animal groups (CTL vs. PAB) denoted by (*) for $p < 0.05$ and (**) for $p < 0.01$.

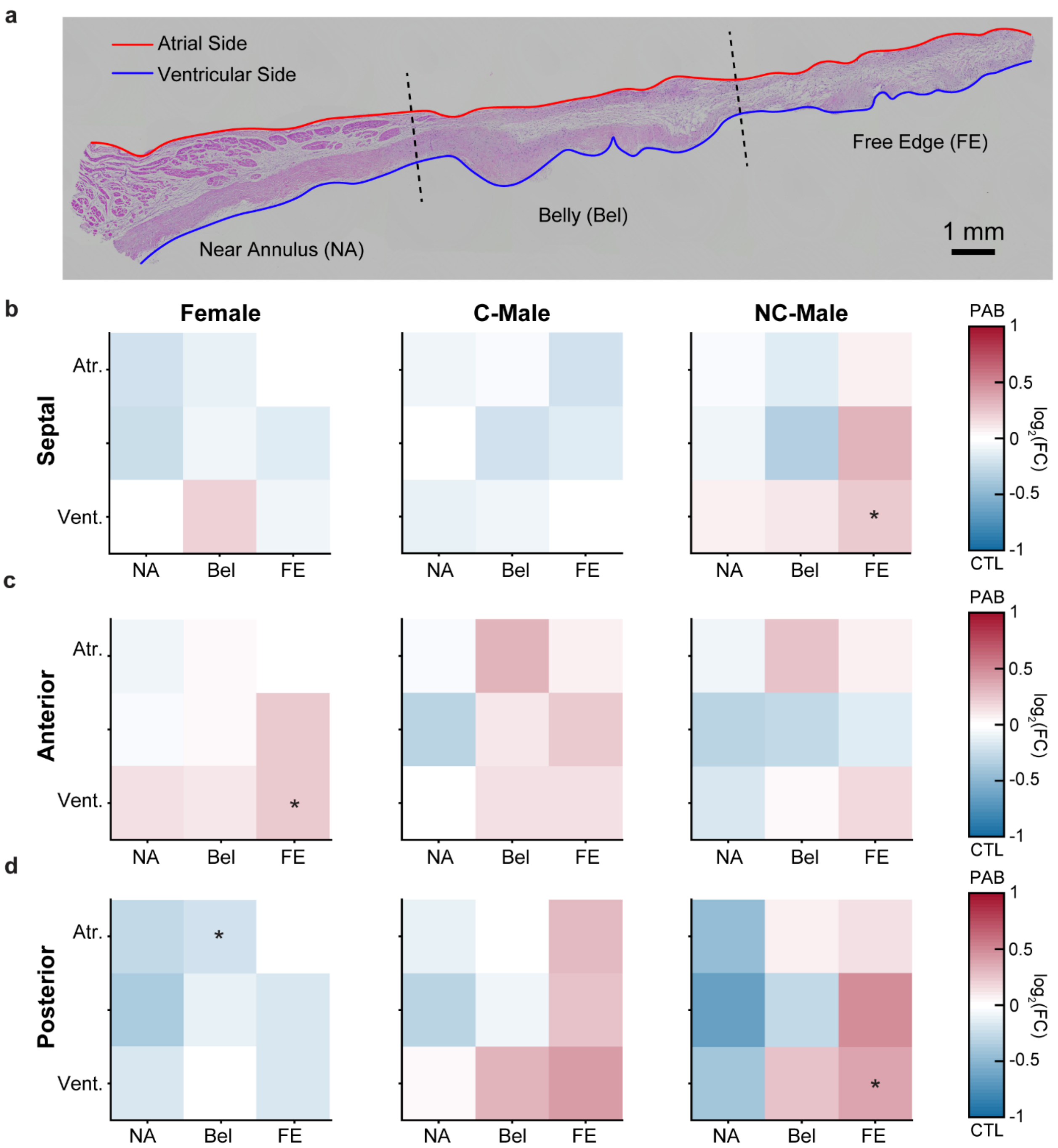


**Figure S11.** Regional changes in leaflet cell nuclei density after PAB are most pronounced near the ventricular free edge and differ by sex and leaflet. (a) Representative radial strip from an anterior leaflet of a PAB sheep showing the near-annulus (NA), belly (Bel), and free-edge (FE) regions, with splines drawn along the atrial (Atr., red) and ventricular (Vent., blue) surfaces. Scale bar = 1 mm. Regional cell nuclei density comparisons for the (b) septal, (c) anterior, and (d) posterior leaflets in female, castrated male (C-Male), and non-castrated male (NC-Male) sheep. Fold change (FC) between PAB and CTL was calculated as the ratio of nuclei per unit area for corresponding leaflet regions. The heat-map color scale reports the logarithm base 2 of the FC, interpreted as (positive, red): PAB leaflet nuclei density is higher than CTL, (0, white): PAB and CTL leaflet nuclei densities are approximately equal, and (negative, blue): PAB leaflet nuclei density is less than CTL. Regional statistical differences between groups were assessed using linear mixed-effects models with post hoc pairwise comparisons of estimated marginal means. Significant contrasts between treatment groups (CTL vs. PAB) are denoted by (*) for $p < 0.05$.

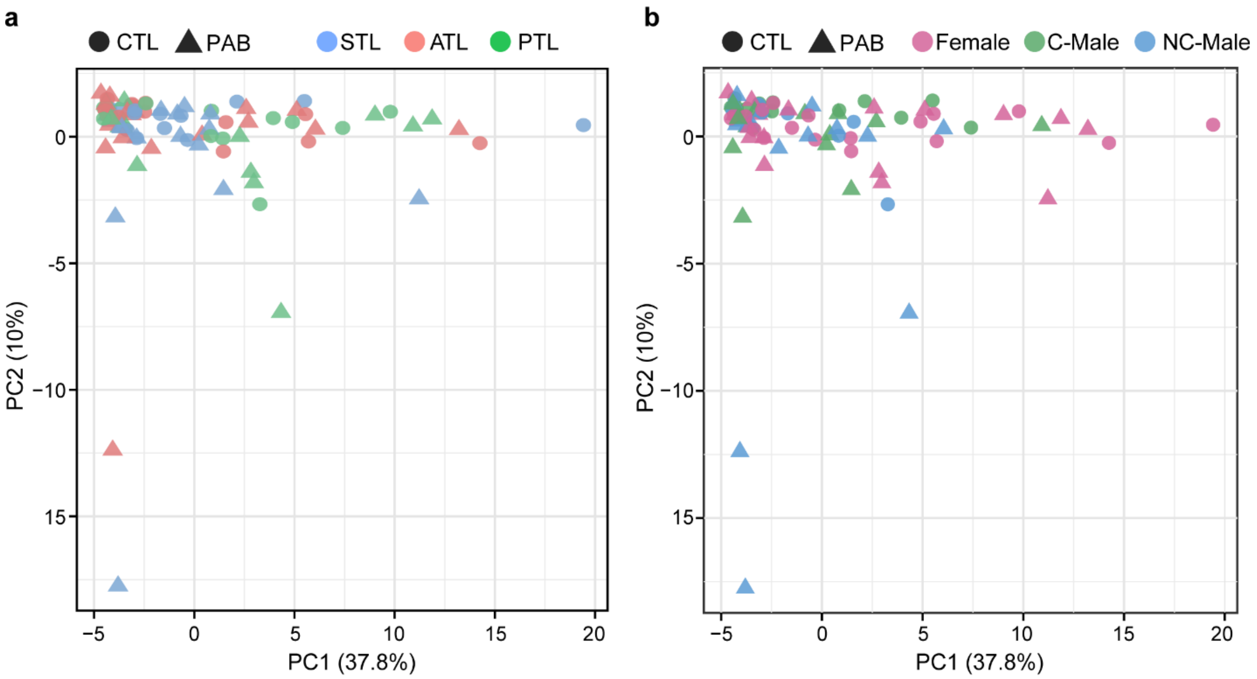


**Figure S12.** Principal component analysis (PCA) of tricuspid leaflet transcriptomes by sex, condition, and leaflet. PCA of regularized log-transformed counts of the top 500 most variable genes for control (CTL) and pulmonary artery banding (PAB) sheep. (a) Each point represents one septal (STL), anterior (ATL), and posterior (PTL) tricuspid leaflet and (b) each point represents one leaflet sample stratified by sex (female, castrated male (C-Male), or non-castrated male (NC-Male)).

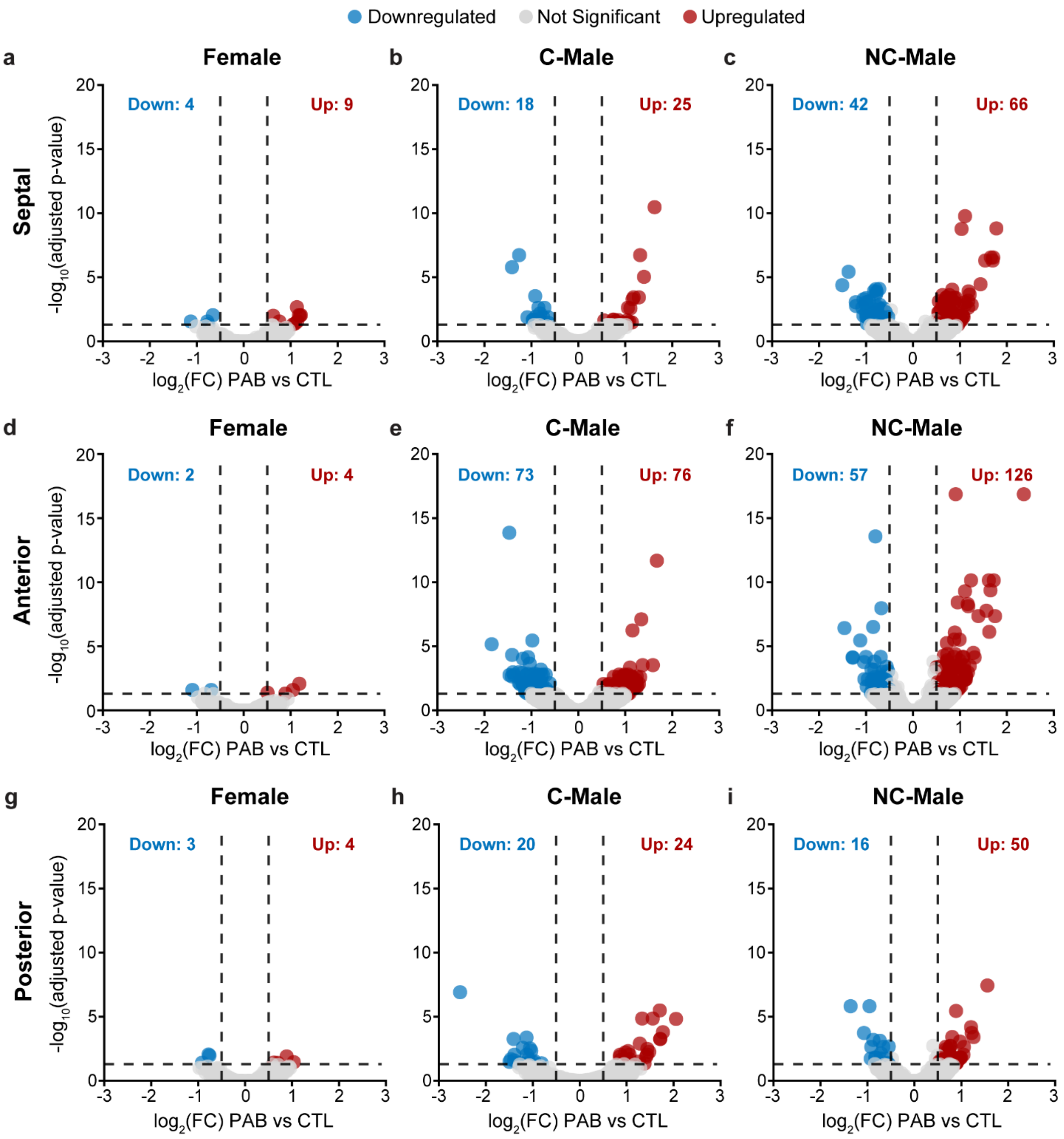


**Figure S13**. Leaflet- and sex-stratified differential gene expression in tricuspid valve leaflets after pulmonary artery banding (PAB). The volcano plots depict differential gene expression comparisons (PAB vs. control [CTL]) of (a-c) septal, (d-f) anterior, and (g-i) posterior tricuspid valve leaflets in Females, castrated males (C-Males), and non-castrated males (NC-Males). Each point represents a gene, plotted along log2-fold change (FC) and –log10(adjusted p-value). Significantly upregulated and downregulated genes (|log2FC| > 0.5 and adjusted p < 0.05) are shown in red and blue, respectively; non-significant genes are shown in grey. Dashed lines indicate significance thresholds. The total number of significantly upregulated and downregulated genes are reported in each panel.

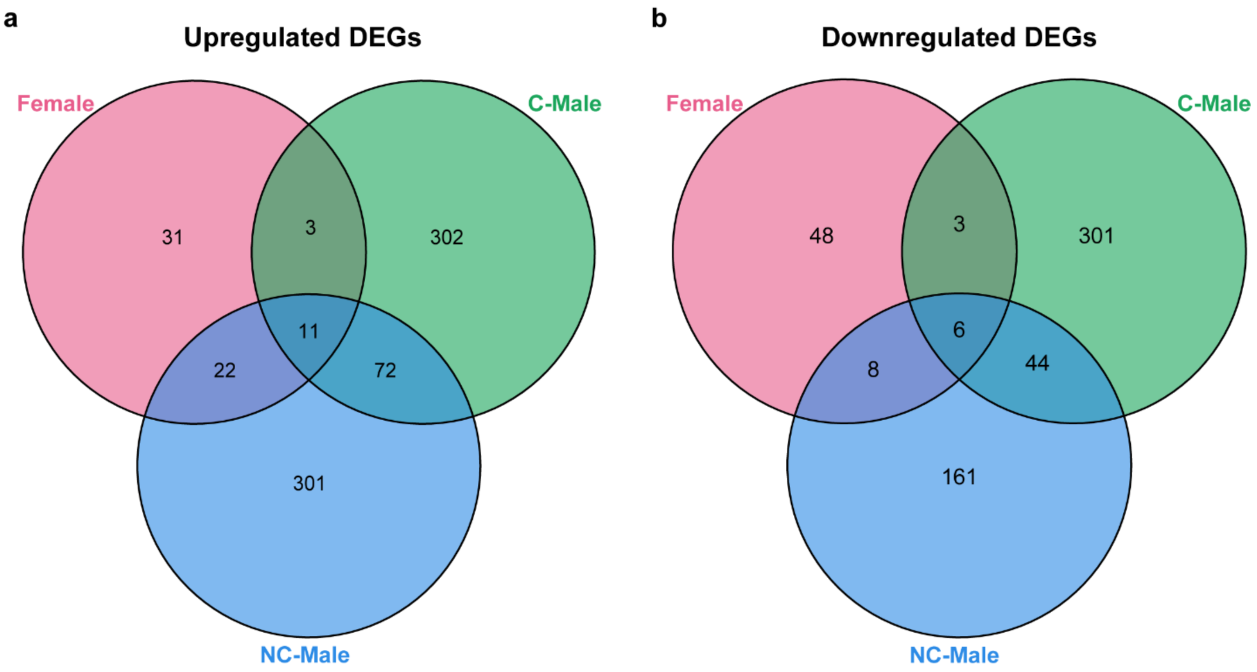


**Figure S14.** Overlap of sex-stratified differentially expressed genes (DEGs) after pulmonary artery banding. (a) Venn diagram depicting overlap of upregulated DEGs in female, castrated male (C-Male), and non-castrated male (NC-Male) tricuspid leaflets following PAB versus CTL (adjusted $p < 0.05$, log2 fold-change $> 0.5$). (b) Venn diagram depicting overlap of downregulated DEGs in the same sex groups (adjusted $p < 0.05$, log2 fold-change $< 0.5$).

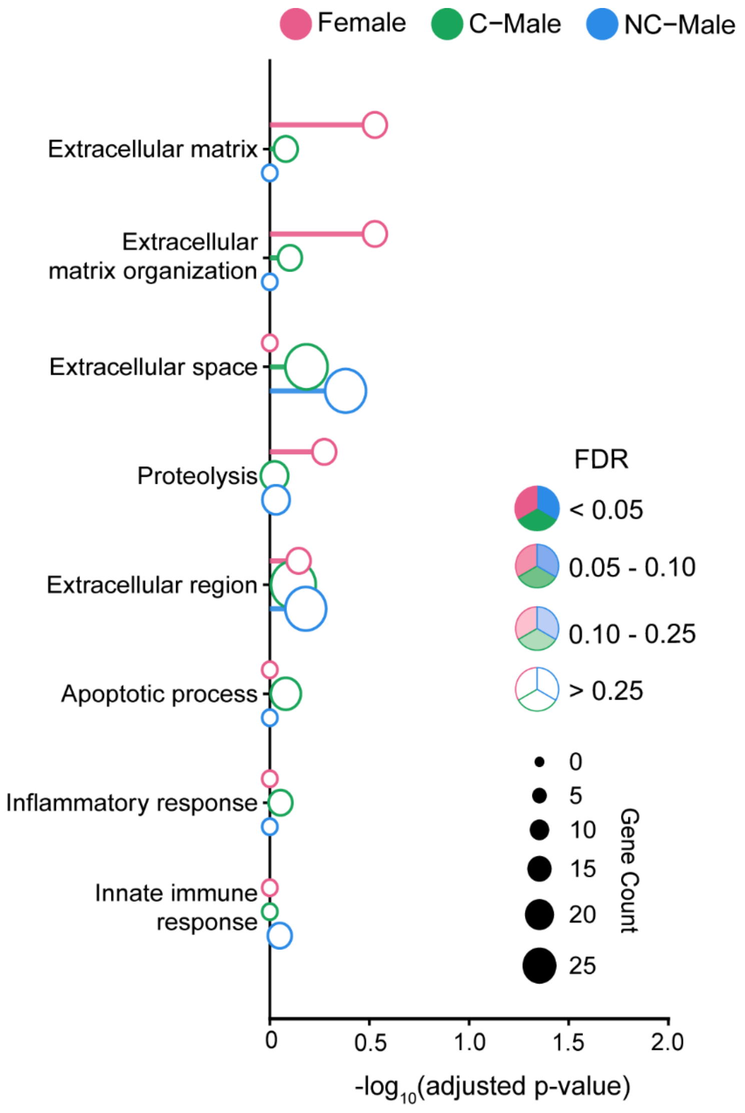


**Figure S15**. Curated Gene Ontology (GO) biological process enrichment analysis of downregulated genes within each sex-stratified comparison. The panel focuses on ECM-, proteolysis-, apoptosis-, and immune-associated pathways. Each point represents a sex-specific enrichment result for a given pathway; point size reflects gene count and the x-axis denotes –log10(adjusted p-value). Pathways meeting FDR < 0.1 are considered significantly enriched.
